# Prediction and preview strongly affect reading times but not skipping during natural reading

**DOI:** 10.1101/2021.10.06.463362

**Authors:** Micha Heilbron, Jorie van Haren, Peter Hagoort, Floris P. de Lange

## Abstract

In a typical text, readers look much longer at some words than at others and fixate some words multiple times, while skipping others altogether. Historically, researchers explained this variation via low-level visual or oculomotor factors, but today it is primarily explained in terms of cognitive factors, such as how well word identity can be predicted from context or discerned from parafoveal preview. While the existence of these effects has been well established in experiments, the relative importance of prediction, preview and low-level factors for eye movement variation in natural reading is unclear. Here, we address this question in three large datasets (n=104, 1.5 million words), using a deep neural network and Bayesian ideal observer to model linguistic prediction and parafoveal preview from moment to moment in natural reading. Strikingly, neither prediction nor preview was important for explaining word skipping – the vast majority of skipping was explained by a simple oculomotor model. For reading times, by contrast, we found strong but independent contributions of both prediction and preview, with effect sizes matching those from controlled experiments. Together, these results challenge dominant models of eye movements in reading by showing that linguistic prediction and parafoveal preview are not important determinants of word skipping.

## Introduction

When reading a text, readers move their eyes across the page to bring new information to the centre of the visual field, where perceptual sensitivity is highest. While it may subjectively feel as if the eyes smoothly slide along the text, they in fact traverse the words with rapid jerky movements called *saccades*, followed by brief stationary periods called *fixations*. Across a text, saccades and fixations are highly variable and seemingly erratic: Some fixations last less than 100 ms, others more than 400; and while some words are fixated multiple times, many other words are skipped altogether [1, 2]. What explains this striking variation?

Historically, researchers have pointed to low-level non-linguistic factors like word length, oculomotor noise, or the relative position where the eyes happen to land [2–5]. Such explanations were motivated by the idea that oculomotor control was largely *autonomous*. In this view, readers can adjust saccade lengths and fixation durations to global characteristics like text difficulty or reading strategy, but not to subtle word-by-word differences in language processing [2–4, 6].

As reading was studied in more detail, however, it became clear that the link between eye movements and cognition was more direct. For instance, it was found that fixation durations were shorter for words with higher frequency [7, 8]. Eye movements were even shown to depend on how well a word’s identity could be inferred *before* fixation. Specifically, researchers found that words are read faster and skipped more often if they are *predictable* from linguistic context [9, 10] or if they are identifiable from a *parafoveal preview* [11–13].

These demonstrations of a direct link between eye movements and language processing overturned the autonomous view, replacing it by cognitive accounts describing eye movements during reading as largely, if not entirely, controlled by linguistic processing [14, 15]. Today, many studies still build on the powerful techniques like gaze-contingent displays that helped overturn the autonomous view, but now ask much more detailed questions, like whether word identification is a distributed or sequential process [16, 17]; how many words can be processed in the parafovea [18]; at which level they are analysed [19, 20], and how this may differ between writing systems or orthographies [21, 22].

Here, we ask a different, perhaps more elemental question: how much of the variation in eye movements do linguistic prediction, parafoveal preview, and non-linguistic factors each explain? That is, how important are these factors for determining how the eyes move during reading? Dominant, cognitive models explain eye movement variation primarily as a function of on-going processing. Skipping, for instance, is modelled as the probability that a word is identified before fixation [14, 23, 24]. Some, however, have questioned this purely cognitive view, suggesting that low-level features like word eccentricity or length might be more important [25–27]. Similarly, one may ask what drives next-word identification: is identifying the next word mostly driven by linguistic predictions [28] or by parafoveal perception? Remarkably, while it is well-established that both linguistic and oculomotor, and both predictive and parafoveal processing, all affect eye-movements [13, 25, 29, 30], a comprehensive picture of the their relative explanatory power is currently missing, perhaps because they are seldom studied all at the same time.

To arrive at such a comprehensive picture we focus on natural reading, analysing three large datasets of participants reading passages, long articles, and even an entire novel – together encompassing 1.5 million (un)fixated words, across 108 individuals [31–33]. Instead of manipulating word predictability or perturbing parafoveal perceptibility, we combine deep neural language modelling [34] and Bayesian ideal observer analysis [35] to quantify how much information is conveyed by both factors, on a moment-by-moment basis. This way, we can probe the effect of both prediction and preview on *each* word during natural reading. Such a broad-coverage approach has been applied to the effects of predictability on reading before [30, 36–39], but either without considering preview or only through coarse heuristics such as using frequency as a proxy for parafoveal identifiability [17, 40, 41] (cf. [35]). By contrast, here we explicitly model both, in addition to low-level explanations like autonomous oculomotor control. To assess explanatory power, we use set theory to derive the unique and shared variation in eye movements explained by each model.

To preview the results, this revealed a striking dissociation between skipping and reading times. For word skipping, the overwhelming majority of variation could be explained – mostly *uniquely* explained – by a non-linguistic oculomotor model. For reading times, by contrast, we found strong effects of both prediction and preview, tightly matching effect sizes from controlled designs. Interestingly, linguistic prediction and parafoveal preview seem to operate independently: we found strong evidence against Bayes-optimal integration of the two. Together, these results challenge dominant cognitive models of reading, and show that skipping (or the decision of *where* to fixate) and reading times (i.e. *how long* to fixate) are governed by different principles.

## Results

We analysed eye movements from three large datasets of participants reading texts ranging from isolated paragraphs to an entire novel. Specifically, we considered three datasets: Dundee [33] (N=10, 51.502 words per participant), Geco [32] (N=14, 54.364 words per participant) and Provo [31] (N=84, 2.689 words per participant). In each corpus, we analysed both skipping and reading times (indexed by gaze duration), as they are thought to reflect separate processes: the decision of *where* vs *how long* to fixate, respectively [14, 25]. For more descriptive details about the data across participants and datasets, see *Methods* and Figures S5-S7.

To estimate the effect of linguistic prediction and parafoveal preview, we quantified the amount of information conveyed by both factors for each word in the corpus (for preview, this was tailored to each individual participant, since each word was previewed at a different eccentricity by each participant). To this end, we formalised both processes as a probabilistic belief about the identity of the next word, given either the preceding words (prediction) or a noisy parafoveal percept (preview; see Figure 1a). As such, we could describe these disparate cognitive processes using a common information-theoretic currency. To compute the probability distributions, we used GPT-2 for prediction [34] and a Bayesian ideal observer for preview [35] (see Figure 1b and *Methods*).

**Figure 1:**
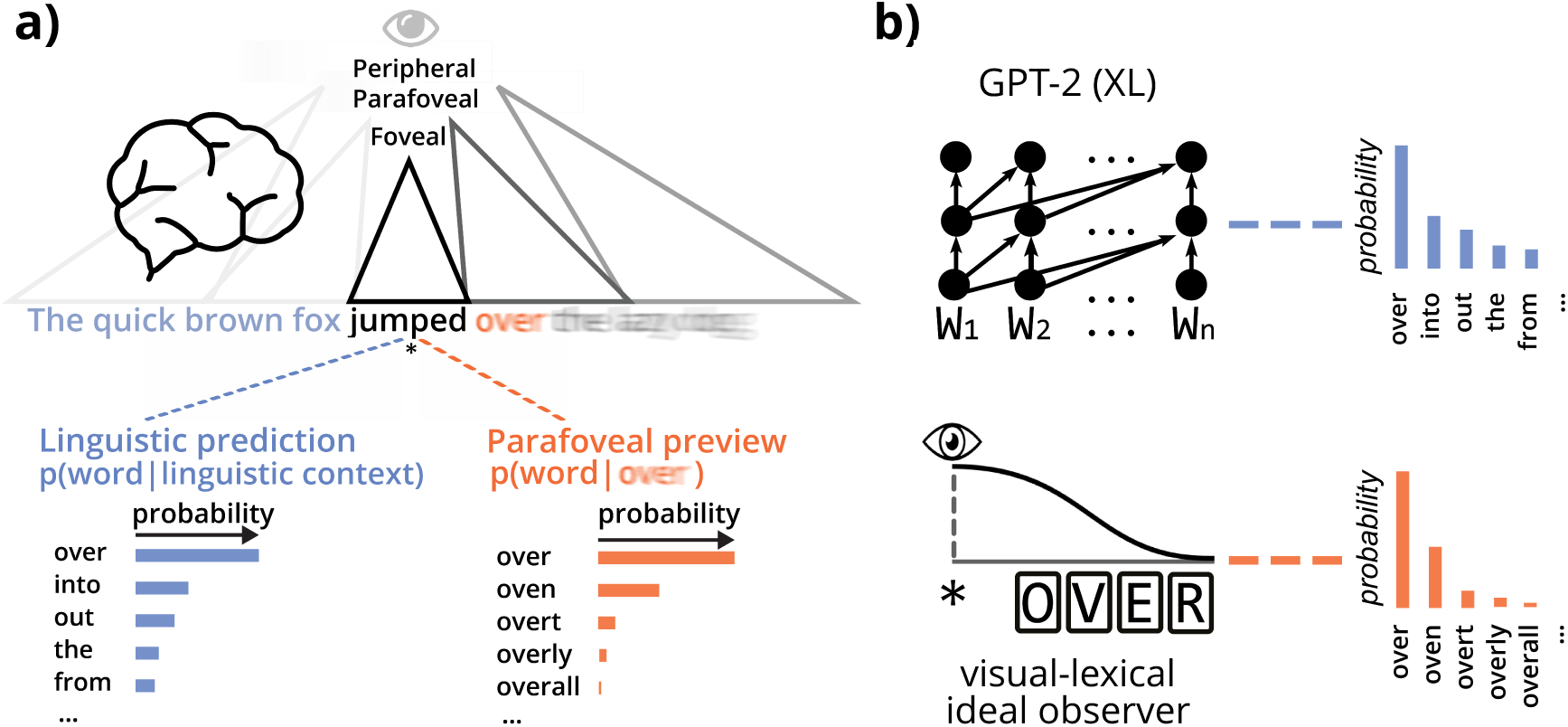
Quantifying two types of context during natural reading. **a)** Readers can infer the identity of the next word before fixation either by predicting it from context or by discerning it from the parafovea. Both can be cast as a probabilistic inference about the next word, either given the preceding words (prediction, blue) or given a parafoveal percept (preview, orange). **b)** To model prediction, we use GPT-2, one of the most powerful publicly available language models [34]. For preview, we use an ideal observer [35] based on well-established ‘Bayesian Reader’ models [42–44]. Importantly, we do not use either model as a cognitive model *per se*, but rather as a tool to quantify how much information is *in principle* available from prediction or preview on a moment-by-moment basis.

### Prediction and preview increase skipping rates and reduce reading times

We first asked whether our formalisations allowed us to observe the expected effects of prediction and preview, while statistically controlling for oculomotor and lexical variables in a multiple regression model. Because the decisions of whether to skip and how long to fixate a word are made at different moments, we modeled each separately with a different set of explanatory variables; but for both, we considered the full model (detailed below).

As expected, we found in all datasets that words were more likely to be skipped if there was more information available from the linguistic prediction (Bootstrap: Dundee, *p* = 0.023; GECO, *p* = 0.034; Provo *p <* 10^−5^) and/or the parafoveal preview (Bootstrap: Dundee, *p* = 4 × 10^−5^; GECO, *p <* 10^−5^; Provo *p <* 10^−5^). Similarly, reading times were reduced for words that were more predictable (all *p*′*s <* 3.2 × 10^−4^) or more identifiable from the parafovea (all *p*′*s <* 4 × 10^−5^).

Together this confirms that our model-based approach can capture the expected effects of both prediction [15] and preview [13] in natural reading, while statistically controlling for other variables.

### Skipping can be largely explained by non-linguistic oculomotor factors

After confirming that prediction and preview had a statistically significant influence on word skipping and reading times, we went on to assess their relative explanatory power. That is, we asked the question how important these factors were, by examining how much variance was explained by each. To this end, we grouped the variables from the full regression model into different types of explanations, and assessed how well each type accounted for the data.

For skipping, we considered three explanations. First, a word might be skipped *purely* because it could be predicted from context – i.e. purely as a function of the amount of information conveyed by the prediction. Secondly, a word might be skipped because its identity could be gleaned from a parafoveal preview – that is, purely as a function of the informativeness of the preview. Finally, a word might be skipped simply because it is so short or so close to the prior fixation location that an autonomously generated saccade will likely overshoot it, irrespective of its linguistic properties – in other words, purely as a function of length and eccentricity. Note that we did not include often-used lexical attributes like frequency to predict skipping, because using attributes of word_*n*+1_ already presupposes parafoveal identification. Moreover, to the extent that a lexical attribute like frequency might influence a words parafoveal identifiability, this should already be captured by the parafoveal entropy (see Figure S3 and *Methods* for more details).

For each word, we thus modelled the probability of skipping either as a function of prediction, preview, or oculomotor information, or by any combination of the three. Then we partitioned the unique and shared cross-validated variation explained by each account. Strikingly, this revealed that the overwhelming majority of explained skipping variation (94 %) could be accounted for by the non-linguistic baseline (Figure 2). Moreover, the majority of the variation was *only* explained by the baseline, which explained 10 times more unique variation than prediction and preview combined. There was a large degree of overlap between preview and the oculomotor baseline, which is unsurprising since a word’s identifiability decreases as a function of its eccentricity and length. Interestingly, there was even more overlap between the prediction and baseline model: almost all skipping variation that could be explained by contextual constraint could be equally well explained by the oculomotor baseline factors.

**Figure 2:**
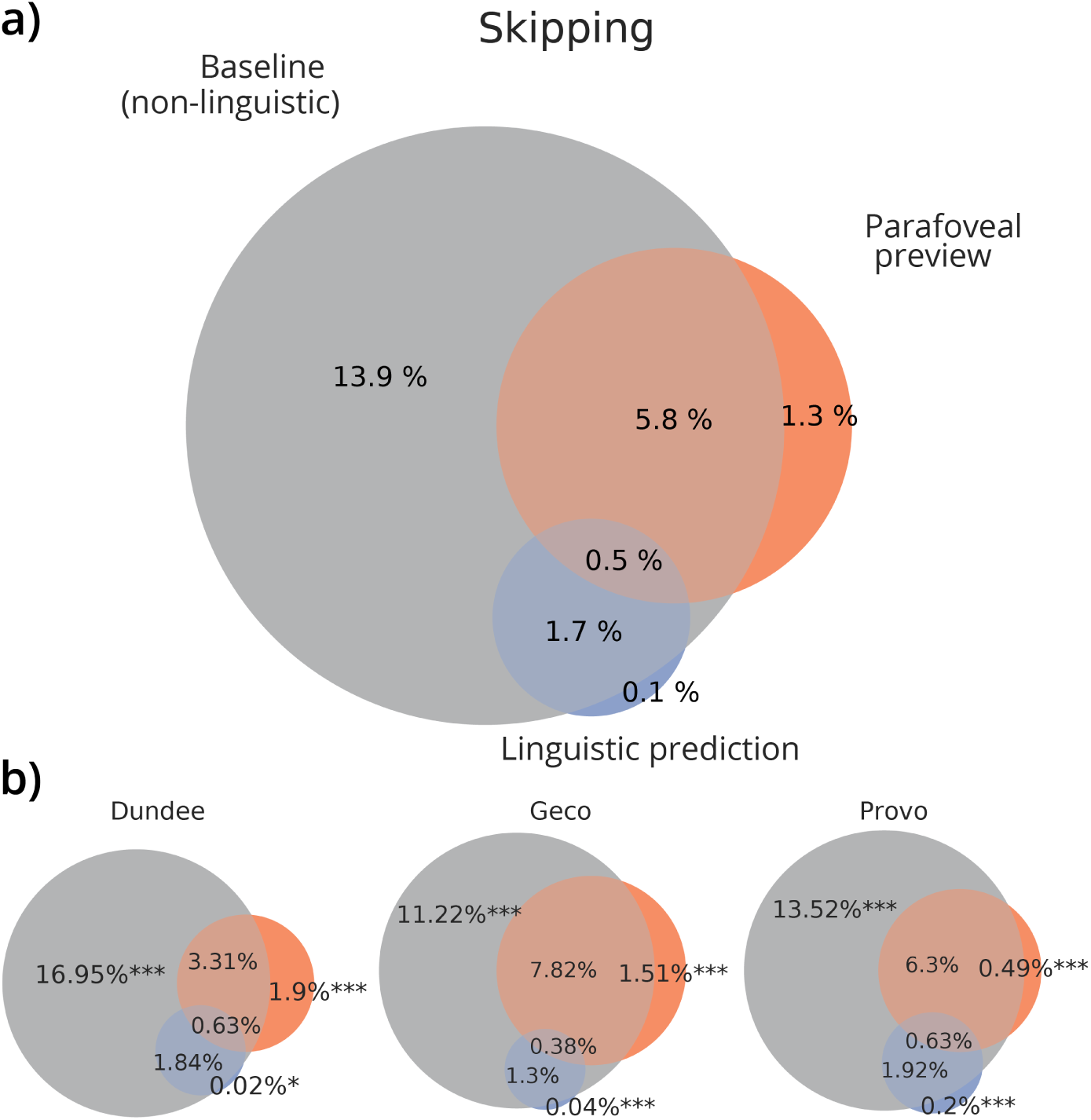
Variation in skipping explained by predictive, parafoveal and autonomous oculomotor processing. **a)** Proportions of cross-validated variation explained by prediction (blue), preview (orange) oculomotor baseline (grey) and their overlap; averaged across datasets (each dataset weighted equally). **b)** Variation partitions for each individual dataset, including statistical significance of variation uniquely explained by predictive, parafoveal or oculomotor processing. Stars indicate significance-levels of the cross-validated unique variation explained (bootstrap t-test against zero): *p <* 0.05 (*), *p <* 0.05 (**), *p <* 0.001 (***) For results of individual participants, and their consistency, see Figure S9.

Importantly, while the contribution of prediction and preview was small, it was significant both for prediction (bootstrap: Dundee, *p* = 0.015; Geco, *p* = 0.0001; Provo, *p <* 10^−5^) and preview (all *p*′*s <* 5 × 10^−5^), confirming that both factors do affect skipping. Crucially however, the vast majority of skipping that could be explained by either prediction or preview was equally well explained by the more parsimonious oculomotor model – which also explained much more of the skipping data overall. This challenges the idea that word identification is the driver behind skiping, instead pointing to a simpler strategy, primarily based on length and eccentricity.

What might this simpler strategy be? One possibility is a ‘blind’ random walk: generating saccades of some average length, plus oculomotor noise. However, we find that saccades are tailored to word length and exhibit a well-known preferred landing position, slightly left to a word’s center (see Figure S8; compare [45, 46]). This suggests the decision of where to look next is not ‘blind’ but based on a coarse low-level visual analysis of the parafovea, for instance conveying just the location of the next word ‘blob’ within a preferred range (i.e. skipping words too close or short; c.f. [25, 26, 47]). Presumably, such a simple strategy would on average sample visual input conveniently, yielding saccades just large enough for comprehension to keep track. However, if such an ‘autopilot’ is indeed independent, one would expect it occasionally go out of step, such that a skipped word cannot be recognised or guessed, derailing comprehension. In line with this, we find evidence for a compensation strategy. The probability that initially skipped word are subsequently (regressively) fixated is strongly, inversely related to its parafoveal identifiability *before* skipping (see Figure S10; logistic regression bootstrap test, all *p*’s *<* 10^−5^). Together, this suggests that initial skipping decisions are primarily driven by a low-level oculomotor ‘autopilot’, but one that is kept in line with online language comprehension by directly correcting saccades that outrun recognition.

### Reading times are strongly modulated by prediction and preview

For reading times (operationalised through gaze durations, so considering foveal ‘reading’ only), we also considered three explanatory factors. First, a word might be read faster because it was predictable from the preceding context, which we formalised via linguistic surprise. Second, a word might be read faster if it could already be partly identified from the parafoveal preview (before fixation). This informativeness of the preview was again formalised via the parafoveal preview entropy. Finally, a word might be read faster due to attributes of the word itself, such as lexical frequency. This last explanatory factor functioned as a baseline that captured key non-contextual word attributes, both linguistic and non-linguistic (see Methods).

In all datasets, prediction (all *p*′*s <* 7.7 × 10^−3^), preview (all *p*′*s <* 1.2 × 10^−4^) and non-contextual woord attributes (all *p*′*s <* 1.8 × 10^−4^) again all explained significant unique variation. The non-contextual baseline explained the most variance, which shows – perhaps unsurprisingly – that properties of the word itself are more important than contextual factors in determining how long a word is fixated. Critically however, compared to skipping the *unique* contribution of prediction and preview was more than three times higher (see Fig 3). Specifically, while prediction and preview could only uniquely account for 6% of explained word skipping variation, they uniquely accounted for more than 18 % of explained variation in reading times. Importantly, the *non-contextual* baseline used to predict reading times included both linguistic (e.g. lexical frequency) and non-linguistic information (viewing position) of the current word. When we analysed these separately, we found that the unique contribution of non-linguistic factors was small (see Figure S12). This shows that contrary to skipping, variation in reading time is heavily influenced by online linguistic processing.

**Figure 3:**
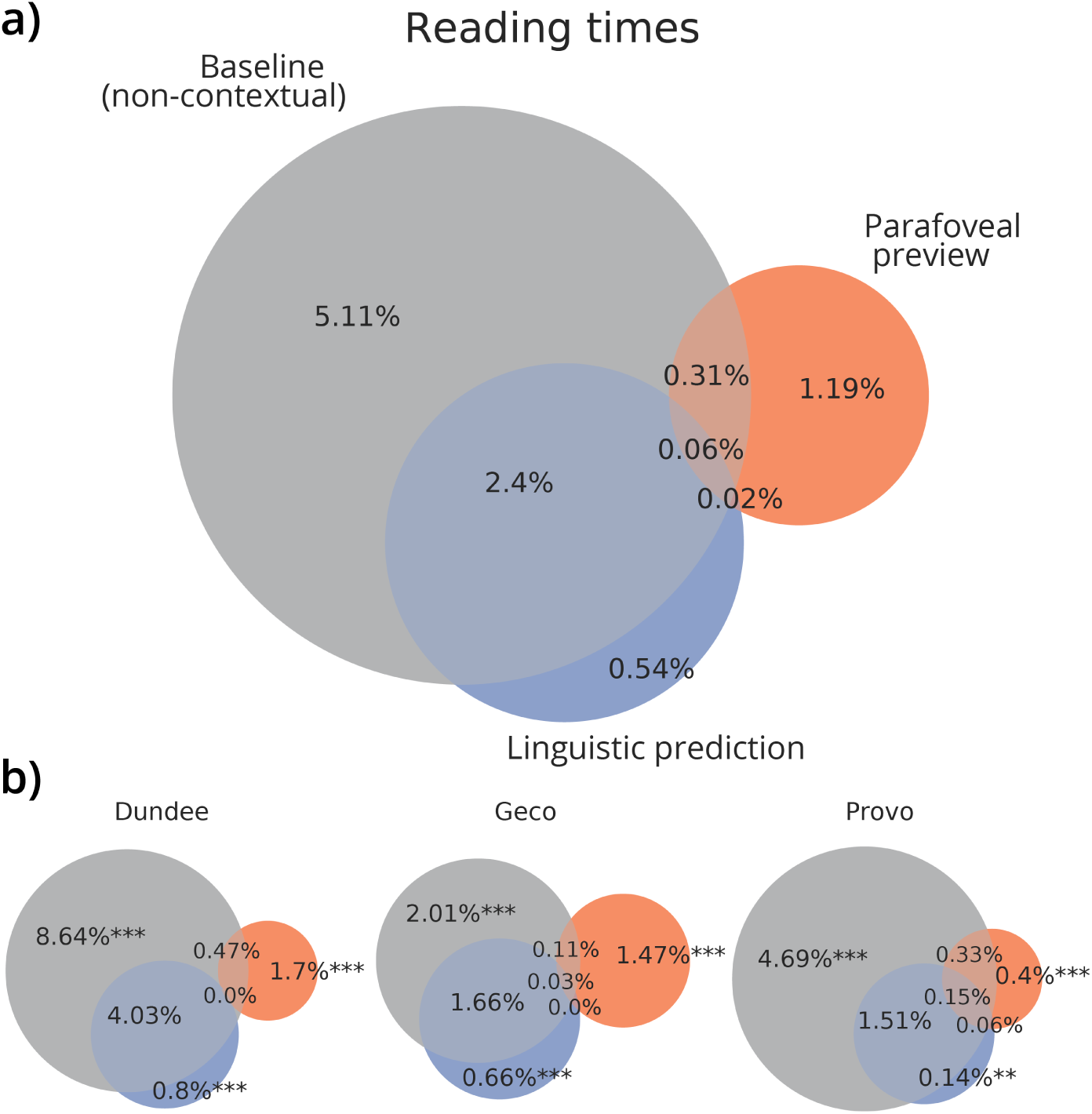
Variation in reading times explained by predictive, parafoveal and non-contextual information. **a)** Grand average of partitions of cross-validated variance in reading times (indexed by gaze durations) across datasets (each dataset weighted equally) explained by non-contextual factors (grey), parafoveal preview (orange), and linguistic prediction (blue). **b)** Variance partitions for each individual dataset, including statistical significance of the cross-validated variance explained uniquely by the predictive, parafoveal or non-contextual explanatory variables. Stars indicate significance-levels of the cross-validated unique variance explained (bootstrap t-test against zero): *p <* 0.05 (**), *p <* 0.001 (***). Note that the non-contextual model here included both lexical attributes (e.g. frequency) and oculomotor factors (relative viewing or landing position); assessing these separately reveals that reading time variation uniquely explained by oculomotor factors was small (see Fig S12). For results of individual participants, see Figure S11.

### Naturalistic prediction and preview benefit effect match experimental effect sizes

The previous result confirms that reading times (indexed via gaze durations) are highly sensitive to linguistic and parafoveal context, in line with decades of research on eye movements in reading [48]. But how well do our results compare exactly to established findings from the experimental literature?

To directly address this question, we simulated, for each participant the effect size of two well-established effects that would be expected to be obtained if we would conduct a well-controlled factorial experiment. Specifically, because we estimated how much additional information from either prediction or preview (in bits) reduced reading times (in milliseconds) we could predict reading times for words that are expected vs unexpected (predictability benefit [29, 49]) or have valid vs invalid preview (i.e. preview benefit [13]).

The simulated effects tightly corresponded to those from experimental studies (see Fig 4). This shows that our analysis does not underfit or otherwise underestimate the effect of prediction or preview. Moreover, it shows that the effect sizes, which are well-established in controlled designs, generalise to natural reading. This is especially interesting for the preview benefit, because it implies that the effect can be largely explained in terms of parafoveal lexical identification [20, 48], and that other factors such as low-level visual ‘preprocessing’, or interference between the (invalid) parafoveal percept and foveal percept, may only play a minor role [c.f. 13, 14].

**Figure 4:**
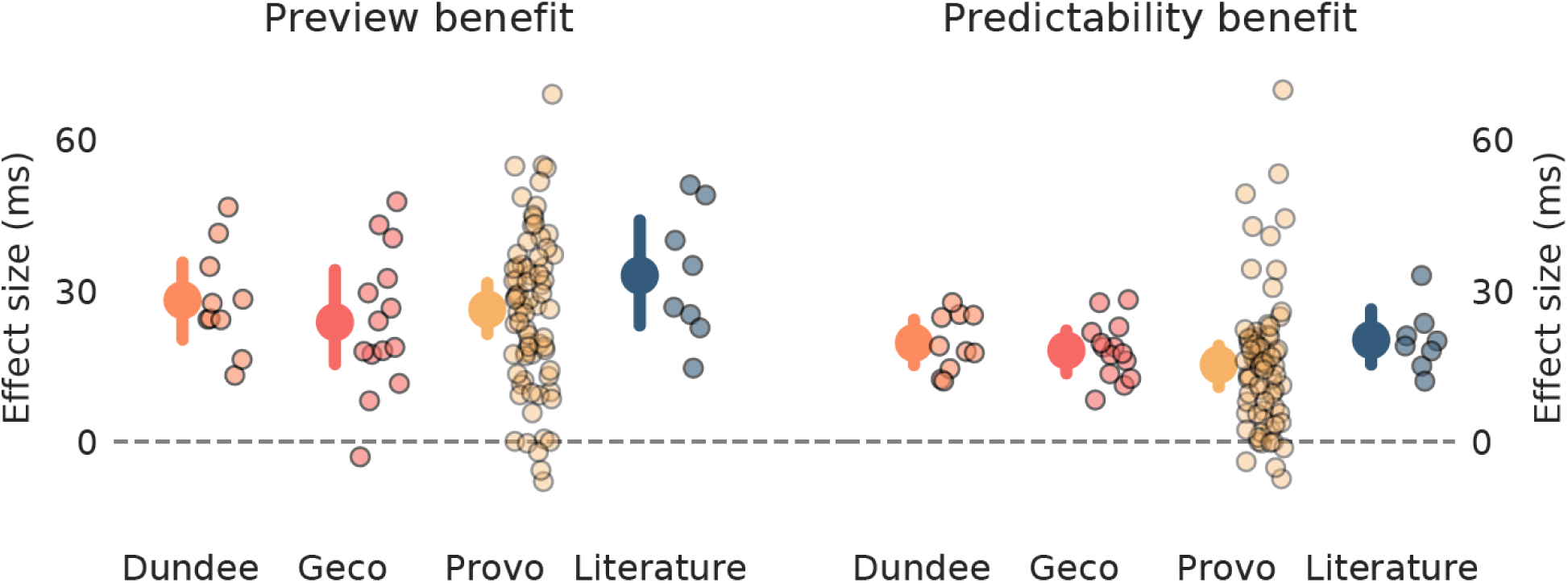
Simulated preview and predictability benefits match those reported in experimental literature. Preview (left) and predictability benefits (right) inferred from our analysis of each dataset, and observed in a sample of studies (see Table S1). In this analysis, preview benefit was simulated as the expected difference in gaze duration after a preview of average informativeness versus after no preview at all. Predictability benefit was defined as the difference in gaze duration for high versus low probability words; ‘high’ and ‘low’ were defined by subdividing the cloze probabilities from provo into equal thirds of ‘low’, ‘medium’ and ‘high’ probability (see Methods). In each plot, small dots with dark edges represent either individual subjects within one dataset or individual studies in the sample of the literature; larger dots with error bars represent the mean effect across individuals or studies, plus the bootstrapped 99% confidence interval.

### No integration of prediction and preview

So far, we have treated prediction and preview as being independent. However, it might be that these processes, while using different information, are integrated – such that a word is parafoveally more identifiable when it is *also* more predictable in context. Bayesian probability theory proposes an elegant and mathematically optimal way to integrate these sources of information: the prediction of the next word could be incorporated as a prior in perceptual inference. Such a contextual prior fits into hierarchical Bayesian models of vision [50], and has been observed in speech perception, where a contextual prior guides the recognition of words from a partial sequence of phonemes [51]. Does such a prior also guide word recognition in reading, based on a partial parafoveal percept?

To test this, we recomputed the parafoveal identifiability of each word for each participant, but now with an ideal observer using the prediction from GPT-2 as a prior. As expected, bayesian integration enhanced perceptual inference: on average, the observer using linguistic prediction as a prior extracted more information from the preview (± 6.25 bits) than the observer not taking the prediction into account (± 4.30 bits; 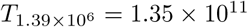, *p* ≈ 0). Interestingly however, it provided a worse fit to the human reading data. This was established by comparing two versions of the full regression model: one with parafoveal entropy from the (theoretically superior) contextual ideal observer and one from the non-contextual ideal observer. In all datasets both skipping and reading times were better explained by a model including parafoveal identifiability from the non-contextual observer (skipping: all *p*′*s <* 10^−5^; reading times: *p*′*s <* 10^−5^; see Figure 5).

**Figure 5:**
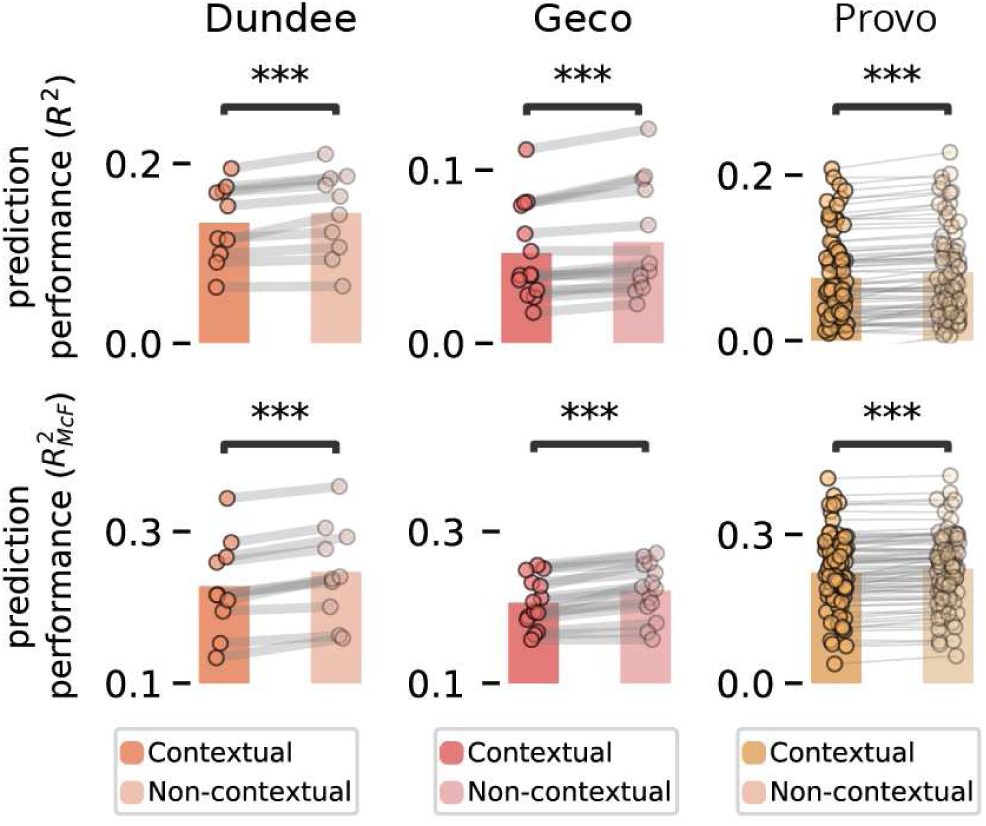
Evidence against bayesian integration of linguistic prediction and parafoveal preview. Cross-validated prediction performance of the full reading times (top) and skipping (bottom) model (including all variables), equipped with parafoveal preview information either from the contextual observer or from the non-contextual observer. Dots with connecting lines indicate participants; stars indicate significance: *p <* 0.001 (***).

Together, this suggests linguistic prediction and parafoveal preview are not integrated, but instead operate independently – thereby highlighting a remarkable sub-optimality in reading.

## Discussion

Eye movements during reading are highly variable. Across three large datasets, we have assessed the relative importance of two major cognitive explanations for this variability – linguistic prediction and parafoveal preview – compared to alternative non-linguistic and non-contextual explanations. This revealed a stark dissociation between skipping and reading times. For word skipping, neither prediction nor preview was especially important, as the overwhelming majority of variation could be explained – mostly *uniquely* explained – by an oculomotor baseline model using just word length and eccentricity. For reading times, by contrast, we observed clear contributions of both prediction and preview (in addition to non-contextual features like frequency) and effect sizes matching those obtained in controlled experiments. Interestingly, preview effects were best captured by a non-contextual observer, suggesting that while readers use both linguistic prediction and preview, these do not appear to be integrated online. Together, the results underscore the dissociation between skipping and reading times, and show that for word skipping, the link between eye movements and cognition is less direct than commonly thought.

Our results on skipping align well with earlier findings by Brysbaert and colleagues [25]. They analysed effect sizes from studies on skipping and found a disproportionately large effect of length, compared to proxies of processing-difficulty like frequency and predictability. We significantly extend their findings by modelling skipping itself (rather than effect sizes from studies) and making a direct link to processing mechanisms. For instance, based on their analysis it was unclear how much of the length effect could be attributed to the lower visibility of longer words – i.e. how much of the length effect may be an identifiability effect [25, p. 19]. We show that length and eccentricity alone explained three times as much variation as parafoveal identifiability – and that most of the variation explained by identifiability was equally well explained by length and eccentricity. This demonstrates that length and eccentricity themselves – not just to the extent they reduce identifiability – are key drivers of skipping.

This conclusion challenges dominant, cognitive models of eye movements, which describe lexical identification as the primary driver behind skipping [14, 23, 24]. Importantly, our results do not challenge predictive or parafoveal word identification itself. Rather, they challenge the notion that moment-to-moment decisions of whether to skip individual words are primarily driven by the recognition of those words. Instead, our results suggest a simpler strategy in which a coarse (e.g. dorsal stream) visual representation is used to reflexively select the next saccade target following the simple heuristic to move forward to the next word ‘blob’ within a certain range (see also [25, 26, 47]). On a neural level, this may imply that saccade target selection is largely independent of word identification in occipitotemporal cortex, and instead relies primarily on a dorsal-frontoparietal visual selection circuit. This circuit can operate separately from higher-order visual analysis, as demonstrated by the fact that visual selection in the frontal eye fields (FEF) can precede object identification in inferior temporal cortex (IT) [52, 53]. During reading, this low-level selection circuit may be the primary determiner of saccadic targets. Influences of identification and comprehension on target selection could then be limited to special cases, such as when the forward drive in FEF is inhibited to make a corrective, backwards saccade.

When conceptualising reading as a process of information sampling [44], such a low-level heuristic for target selection may appear at odds with other accounts of sampling, describing saccade targeting via a drive to reduce uncertainty. These accounts are supported by evidence that saccades are guided to the most informative stimuli [54–56] and that parietal neurons involved in oculomotor decisions encode information gain expected from a saccade, prior to its execution [57, 58]. However, we do not believe that the accounts are strictly at odds, as reading may pose a special case that is quite different from other forms of saccadic sampling. Reading is an over-trained skill in which the space of potential next saccade targets is highly constrained, such that a simple oculomotor strategy may suffice. That said, we do find that the amount of word-identifying information conveyed by preview explains some unique skipping variation [see also 35]. Therefore, it may be that identification-based and oculomotor policies operate at the same time, in a constant competition that the oculomotor heuristic overwhelmingly wins.

Given that readers use both prediction and preview, why would they strongly affect reading times but hardly word skipping? We suggest this is because these different decisions – of *where* versus *how long* to fixate – are largely independent at made at different moments [52, 59, 60]. Specifically, the decision of where to fixate – and hence whether to skip the next word – is made early in saccade programming, which can take 100-150 ms [25, 59, 61]. Although the exact sequence of operations leading to a saccade remains debated, given that readers on average only look some 250 ms at a word, it is clear that skipping decisions are made under strong time constraints, especially given the lower processing rate of parafoveal information. We suggest that the brain meets this constraint by resorting to a computationally frugal ‘move forward’ policy. How long to fixate, by contrast, depends on saccade *initiation*. This process is separate from target selection, as indicated by physiological evidence that variation in target selection time only weakly explains variation in initiation times, which are affected by more factors and can be until adjusted later [52, 60]. This can allow initiation to be informed by foveal information, which is processed more rapidly and may thus more directly influence the decision to either keep dwelling or execute the saccade.

A distinctive feature of our approach is that we focus on a limited number of computationally explicit explanations, rather than using lexical attributes as proxies for explanations (e.g. a word’s frequency as a proxy for its identifiability). For instance, we model preview using a single variable that should capture all effects of variables like frequency on parafoveal identifiability (see Figure S3 and *Methods*). A limitation of this approach is that a model imperfection may prevent one from fully capturing the effect of preview, resulting in an *underestimation*. However, a key advantage of the approach is that it can avoid confound-related *overestimations*. For instance, frequency is correlated with length, so when using frequency as a proxy for parafoveal identifiability, one may find apparent preview effects which are in fact length effects, and strongly overestimate preview importance [62]. Therefore, we only used independent variables that explicitly formalise a cognitive explanation. Based on the effect sizes for gaze duration (Fig 4) we do not believe that this model-based approach significantly underestimates either prediction or preview, and we are optimistic the results provide the comprehensive, interpretable picture we aimed for.

When comparing Figures 2, 3 and 5, the numerical *R*^2^ values of the reading times regression may seem rather small. This could indicate a poor fit, which might undermine our claim that reading times are to a large degree explained by cognitive factors. However, we do not believe this is the case, since our *R*^2^’s for gaze durations are not lower than *R*^2^’s reported by other regression analyses of gaze durations in natural reading [e.g. 17]; and because we find effect sizes in line with the experimental literature (Fig 4). Therefore, we do not believe we overfit or underfit the gaze durations. Instead, what the relatively low *R*^2^ values indicate, we suggest, is that gaze durations are inherently noisy; i.e. that only a limited amount of the variation is *systematic* variation. While this noisiness might be interesting in itself, it is not of interest in this study, which focusses on the relative importance of different *explanations*, and hence only on *systematic* variation. Therefore, what matters is not the absolute *R*^2^ values, but rather the relative importance of different explanations – in other words, the relative size of the circles in Figures 2, 3 and S12, their overlap, and the explanations each circle represents. It is on this level of analysis that we find the stark dissociation – that for skipping (but not for reading times) a simple low-level heuristic can account for almost all of the explained variation – and not on the level of absolute values of variation explained.

A final notable finding is that preview was best explained by a non-contextual observer. This replicates the only other study that compared contextual and non-contextual models of preview [35]. That study focussed on skipping; the fact that we obtain the same result for reading times and in different datasets strengthens the conclusion that prior context does not inform preview. However, this conclusion seems to contradict experiments finding an interaction between context and preview [e.g. 9, 63–65]. One explanation for this discrepancy stems from how the effect is measured. Experimental studies looked at the effect of context on the difference in reading time after valid versus invalid preview [64, 65]. This may reveal a context effect not on recognition, but at a later stage (e.g. priming between context, preview and foveal word). Arguably, these options yield different predictions. If context affects recognition it may allow identification of otherwise unidentifiable words. But if the interaction occurs later it may only *amplify* processing of recognisable words. Constructing a model that formally reconciles this discrepancy is an interesting challenge for future work.

Given that readers use both prediction and preview, why doesn’t contextual prediction *inform* preview? One explanation may be the time constraints imposed by eye movements. Given that readers on average only look some 250 ms at a word in which they have to recognise the foveal word and process the parafoveal percept, this perhaps leaves too little time to fully let the foveal word and context inform parafoveal preview. On the other hand, word recognition based on partial input also occurs in speech perception under significant time-constraints. But despite those constraints, sentence context does influence auditory word recognition [66, 67], a process best modelled by a contextual prior [51, 68]; i.e. the opposite of what we find here. Therefore, rather than being related to time-constraints *per se*, it might be additionally related to the underlying circuitry. More precisely, to the fact that contrary to auditory word recognition, visual word recognition is a laboriously acquired skill that occurs throughout areas in the visual system that are repurposed (rather than evolved) for reading [69, 70]. Therefore, the global sentence context might be able to dynamically influence the recognition of speech sounds in temporal cortex, but not that of words in visual cortex; there, context effects might be confined to simpler, more local context, like lexical context effects on letter perception [71–74].

In conclusion, we have found that two important contextual sources of information in reading, linguistic prediction and parafoveal preview, strongly drive variation in reading times, but hardly affect word skipping, which is largely based on low-level factors. Our results show that as readers, we do not always use all information available to us; and that we are, in a sense, of two minds: consulting complex inferences to decide how long to look at a word, while employing semi-mindless scanning routines to decide where to look next. It is striking that these disparate strategies operate mostly in harmony. Only occasionally they go out of step – then we notice that our eyes have moved too far and we have to look back, back to where our eyes left cognition behind.

## Methods

We analysed eye-tracking data from three, big, naturalistic reading corpora, in which native English speakers read texts while eye-movement data was recorded [32, 33, 39].

### Stimulus materials

We considered the English-native portions of the Dundee, Geco and Provo corpora. The Dundee corpus comprises eye-movements from 10 native speakers from the UK ([33]), who read a total of 56.212 words across 20 long articles from The Independent newspaper. Secondly, the English portion of the Ghent Eye-tracking Corpus (Geco) [32] is a collection of eye movement data from 14 UK English speakers who each read Agathe Cristie’s *The Mysterious Affair at Styles* in full (54.364 words per participant). Lastly, the Provo corpus ([31]) is a collection of eye movement data from 84 US English speakers, who each read a total of 55 paragraphs (extracted from diverse sources) for a total of 2.689 words.

### Eye tracking apparatus and procedure

In all datasets, eye movements were recorded monocularly, by recording the right eye. In Geco and Provo, recordings were made using an EyeLink 1000 (SR Research, Canada) with a spatial resolution of 0.01^°^ and a temporal resolution of a 1000 Hz. For Dundee, a Dr. Bouis oculometer (Dr. Bouis, Kalsruhe, Germany), with a spatial resolution of *<* 0.1^°^ and a temporal resolution of 1000 Hz was used. To minimize head movement, the participant’s heads were stabilised with a chinrest (Geco, Provo) or a bite bar (Dundee). In each experiment, texts were presented in ‘screens’ with either five lines (Dundee) or one paragraph per screen (Geco and Provo), presented using a font size of 0.33^°^ per character. Each screen began with a fixation mark (gaze trigger) that was replaced by the initial word when stable fixation was achieved. In all datasets, a 9-point calibration was performed prior to the recording. In the longer experiments, a recalibration was performed every three screens (Dundee) or either every 10 minutes or whenever the drift correction exceeded 0.5^°^ (Geco) For Dundee and Provo, the order of different texts were randomized across participants. In Geco, the entire novel was read start to finish with breaks between each chapter, during which participants answered comprehension questions.

For each corpus the x,y-values per fixation position were converted into a word-by-word format. In Dundee, raw *x, y*−values were smoothed by rounding to single character precision. In Geco and Provo, raw *x, y*−values for each within-word- or within-letter fixation were preserved and available for each word. Across the three data sets we redefined the bounding boxes around each word, such that they subtended the area between the first to the last character of the word, with the boundary set halfway to the neighboring character (e.g. halfway the before and after the word). Punctuation before or after the word were left out, and words for which the bounding box was inconsistently defined were ignored. For distributions of saccade and fixation data, see Figures S5-S7.

### Language model

Contextual predictions were formalised using a language model – a model computing the probability of each word given the preceding words. Here, we used GPT-2 (XL) – currently among the best publicly released English language models. GPT-2 is a transformer-based model, that in a single pass turns a sequence of tokens (representing either whole words or word-parts) *U* = (*u*_1_, …, *u*_*k*_) into a sequence of conditional probabilities, (*p*(*u*_1_), *p*(*u*_2_|*u*_1_), …, *p*(*u*_*k*_|*u*_1_, …, *u*_*k*−1_)).

Roughly, this happens in three steps: first, an embedding step encodes the sequence of symbolic tokens as a sequence of vectors, which can be seen as the first hidden state *h*_*o*_. Then, a stack of *n* transformer blocks each apply a series of operations resulting in a new set of hidden states *h*_*l*_, for each block *l*. Finally, a (log-)softmax layer is applied to compute (log-)probabilities over target tokens. In other words, the model can be summarised as follows:

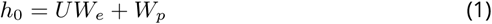

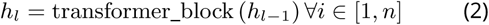

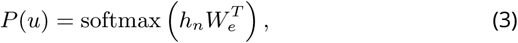

where *W*_*e*_ is the token embedding and *W*_*p*_ is the position embedding.

The key component of the transformer-block is *masked multi-headed self-attention* (Fig S1). This transforms a sequence of input vectors (x_1_, x_2_, … x_*k*_) into a sequence of output vectors (y_1_, y_2_, …, y_*k*_). Fundamentally, each output vector y_*i*_ is simply a weighted average of the input vectors: 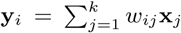. Critically, the weight *w* _*i,j*_ is not a parameter, but is *derived* from a dot product between the input vectors 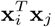, passed through a softmax and scaled by a constant determined by the dimensionality 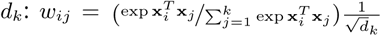. Because this is done for each position, each input vector x_*i*_ is used in three ways: first, to derive the weights for its own output, y_*i*_ (as the *query*); second, to derive the weight for any other output y_*j*_ (as the *key*); finally, in it used in the weighted sum (as the *value*). Different linear transformations are applied to the vectors in each cases, resulting in Query, Key and Value matrices (*Q, K, V*). Putting this all together, we obtain:

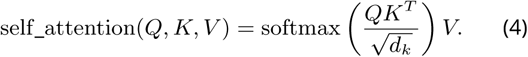

To be used as a language model, two elements are added. First, to make the operation position-sensitive, a position embedding *W*_*p*_ is added during the embedding step – see Equation (1). Second, to enforce that the model only uses information from the past, attention from future vectors is masked out. To give the model more flexibility, each transformer block contains multiple instances (‘heads’) of the self-attention mechanisms from Equation (4).

In total, GPT-2 (XL) contains *n* = 48 blocks, with 12 heads each; a dimensionality of *d* = 1600 and a context window of *k* = 1024, yielding a total 1.5 × 10^9^ parameters. We used the PyTorch implementation of GPT-2 provided by Hugging-Face’s *Transformers* package [75]. For words spanning multiple tokens, we computed their joint probability.

### Ideal observer

To compute parafoveal identifiability, we implemented an ideal observer based on the formalism by Duan & Bicknell [35]. This model formalises parafoveal word identification using Bayesian inference and builds on previous well-established ‘Bayesian Reader’ models [42–44]. It computes the probability of the next word given a noisy percept by combining a prior over possible words with a likelihood of the noisy percept, given a word identity:

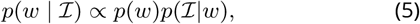

where *ℐ* represents the noisy visual input, and *w* represents a word identity. We considered two priors (see Fig 5): a non-contextual prior (the the overall probability of words in English based on their frequency in Subtlex ([76]), and a contextual prior based on GPT2 (see below). Below we describe how visual information is represented and perceptual inference is performed. For a graphical schematic of the model, see Fig S2; for some distinctive simulations showing how the model captures key effects of linguistic and visual characteristics on word recognition, see Fig S3.

### Sampling visual information

Like in other Bayesian Readers [42–44], noisy visual input is accumulated by sampling from a multivariate Gaussian which is centered on a one-hot ‘true’ letter vector – here represented in an uncased 26-dimensional encoding – with a diagional covariance matrix Σ(*ε*) = *λ*(*ϵ*)^− 1*/*2^*I*. The shape of Σ is thus scaled by the sensory quality *λ*(*ε*) for a letter at eccentricity *ε*. Sensory quality is computed as a function of the perceptual span: this uses using a Gaussian integral based follows the perceptual span or processing rate function from the SWIFT model [23]. Specifically, for a letter at eccentricity *ε, λ* is given by the integral within the bounding box of the letter:

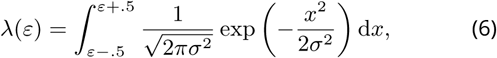

which, following [35, 44], is scaled by a scaling factor Λ. Unlike SWIFT, the Gaussian in Equation 6 is symmetric, since we only perform inference on information about the next word. By using one-hot encoding and a diagonal covariance matrix, the ideal observer ignores similarity structure between letters. This is clearly a simplification, but one with significant computational benefits; moreover, it is a simplification shared by all Bayesian Reader-like models [35, 42, 44], which can nonetheless capture many important aspects of visual word recognition and reading. To determine parameters Λ and *σ*, we performed a grid search on a subset of Dundee and Geco (see Fig S4), resulting in Λ = 1 and *σ* = 3. Note that this *σ* value is close to the average *σ* value of SWIFT and (3.075) and corresponds well to prior literature on the size of the perceptual span (±15 characters; [13, 23, 44]).

### Perceptual inference

Inference is performed over the full vocabulary. This is represented as a matrix which can be seen as a stack of word vectors, y_1_, y_2_, …, y_v_, obtained by concatenating the letter vectors. The vocabulary is thus a *V* × *d* matrix, with *V* the number of words in the vocabulary and *d* the dimensionality of the word vectors (determined by the length of the longest word: *d* = 26 × *l*_*max*_).

To perform inference, we use the belief-updating scheme from [35], in which the posterior at sample *t* is expressed as a (*V* − 1) dimensional log-odds vector x^(t)^, in which each entry 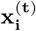 represents the log-odds of y_i_ relative to the final word y_v_. In this formulation, the initial value of x is thus simply the prior log odds, 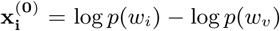, and updating is done by summing prior log-odds and the log-odds likelihood. This procedure is repeated for *T* samples, each time taking the posterior of the previous timestep as the prior in the current timestep. Note that using log odds in this way avoids renormalization:

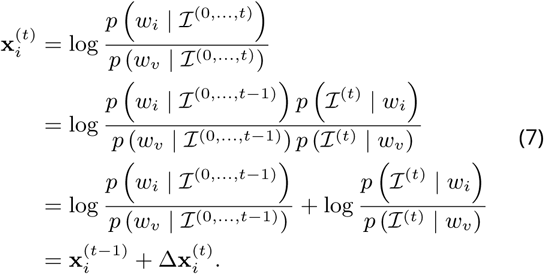

In other words, as visual sample*ℐ* ^(*t*)^ comes in, beliefs are updated by summing the prior log odds x^(*t*−1)^ and the log-odds likelihood of the new information x^(*t*)^.

For a given word *w*_*i*_, the log-odds likelihood of each new sample is the difference of two multivariate Gaussian log likelihoods, one centred on y_*i*_ and one on the last vector y_*v*_. This can be formulated as a linear transformation of *ℐ*:

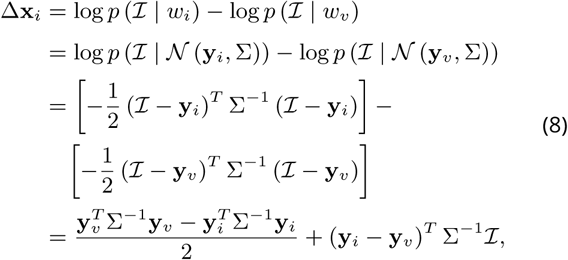

which implies that updating can be implemented by sampling from a multivariate normal. To perform inference on a given word, we performed this sampling scheme until convergence (using *T* = 50), and then transformed the posterior log-odds into the log posterior, from which we computed the Shannon entropy as a metric of parafoveal identifiability.

To compute the parafoveal entropy for each word in the corpus, we make the simplifying assumption that parafoveal preview only occurs during the last fixation prior to a saccade, thus computing the entropy as a function of the word itself and its distance to the last fixation location within the previously fixated word (which is not always the previous word). Because this distance is different for each participant, it was computed separately for each word, for each participant. Moreover, because the inference scheme is based on sampling, we repeated it 3 times, and averaged these to compute the posterior entropy of the word. The amount of information obtained from the preview is then simply the difference between prior and posterior entropy.

The ideal observer was implemented in custom Python code, and can be found in the data sharing collection (see below).

### Contextual vs non-contextual prior

We considered two observers: one with a non-contextual prior capturing the overall probability of a word in a language, and with a contextual prior, capturing the contextual probability of a word in a specific context. For the non-contextual prior, we simply used lexical frequencies from which we computed the (log)-odds prior used in equation (7). For the contextual prior, we derived the contextual prior from log-probabilities from GPT-2. This effectively involves constructing a new Bayesian model for each word, for each participant, in each dataset. To simplify this process, we did not take the full predicted distribution of GPT-2, but only the ‘nucleus’ of the top *k* predicted words with a cumulative probability of 0.95, and truncated the (less reliable) tail of the distribution. Further, we simply assumed that the rest of the tail was ‘flat’ and had a uniform probability. Since the prior odds can be derived from relative frequencies, we can think of the probabilities in the flat tail as having a ‘pseudocount’ of 1. If we similarly express the prior probabilities in the nucleus as implied ‘pseudofrequencies’, the cumulative implied nucleus frequency is then complementary to the the length of the tail, which is simply the difference between the vocabulary size and nucleus size (*V* − *k*). As such, for word *i* in the text, we can express the nucleus as implied frequencies as follows:

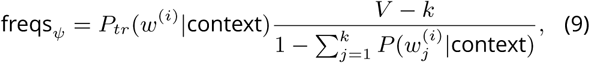

where *P*_*tr*_(*w*^(*i*)^|context) is the trunctated lexical prediction, and 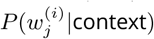 is predicted probability that word *i* in the text is word *j* in the sorted vocabulary. Note that using this flat tail not only simplifies the computation, but also deals with the fact that the vocabulary of GPT-2 is smaller than of the ideal observer – using this tail we can still use the full vocabulary (e.g. to capture orthographic uniqueness effects), while using 95% of the density from GPT-2.

### Data selection

In all our analyses, we focus strictly on first-pass reading, analysing only those fixations or skips when none of the subsequent words have been fixated before. We extensively preprocessed the corpora so that we could include as many words as possible. However, we had to impose some additional restrictions. Specifically we did not include words if they a) contained non-alphabetic characters; b) if they were adjacent to blinks; c) if the distance to the prior fixation location was more than 24 characters (±8^°^); moreover, for the gaze duration we excluded d) words with implausibly short (*<* 70*ms*) or long (*>* 900*ms*) gaze durations. Criterion c) was chosen because some participants occasionally skipped long sequences of words, up to entire lines or more. Such ‘skipping’ – indicated by saccades much larger than the the perceptual span – is clearly different from the skipping of words during normal reading, and was therefore excluded. Note that these criteria are comparatively mild (cf. [35, 36]), and leave approximately 1.1 million observations for the skipping analysis, and 593.000 reading times observations.

### Regression models: skipping

Skipping was modelled via logistic regression in scikit-learn [77], with three sets of explanatory variables (or ‘models’) each formalising a different explanation for why a word might be skipped.

First, a word might be skipped because it could be confidently predicted from context. We formalise this via *linguistic entropy*, quantifying the information conveyed by the prediction from GPT-2. We used entropy, not (log) probability, because using the next word’s probability directly would presuppose that the word is identified, undermining the dissociation of prediction and preview. By contrast, prior entropy specifically probes the information available from prediction only.

Secondly, a word might be skipped because it could be identified from a parafoveal preview. This was formalised via parafoveal entropy, which quantifies the parafoveal preview uncertainty (or, inversely, the amount of information conveyed by the preview). This is a complex function integrating low-level visual (e.g. decreasing visibility as a function of eccentricity) and higher-level information (e.g. frequency or orthographic effects) and their interaction (see Fig S3). Here, too we did not use lexical features (e.g. frequency) of the next word to model skipping directly, as this presupposes that the word is identified; and to the extent that these factors are expected to influence identifiability, this is already captured by the parafoveal entropy (Fig S3).

Finally, a word might be skipped simply because it is too short and/or too close to the prior fixation location, such that a fixation of average length would overshoot the word. This autonomous oculomotor account was formalised by modelling skipping probability purely as a function of a word’s length and its distance to the previous fixation location.

Note that these explanations are not mutually exclusive, so we also evaluated their combinations (see below).

### Regression models: reading time

As an index of reading time, we analysed first-pass *gaze duration*, the sum of a word’s first-pass fixation durations. We analyse gaze durations as they arguably most comprehensively reflect how long a word is looked at, and are the focus of similar model-based analyses of contextual effects in reading [36, 38]. For reading times, we used linear regression, and again considered three sets of explanatory variables, each formalising a different kind of explanation.

First, a word may be read more slowly because it is unexpected in context. We formalised this using surprisal − log(*p*), a metric of a word’s unexpectedness – or how much information is conveyed by a word’s identity in light of a prior expectation about the identity. To capture spillover (R; regpaper; smith) we included not just the surprisal of the current word, but also that of the previous two words.

Secondly, a word might be read more slowly because it was difficult to discern from the parafoveal preview. This was formalised using the parafoveal entropy (see above).

Finally, a word might be read more slowly because of non-contextual factors of the word itself. This is an aggregate baseline explanation, aimed to capture all relevant non-contextual word attributes, which we contrast to the two major contextual sources of information about a word identity that might affect reading times (prediction and preview). We included word class, length, log-frequency, and the relative landing position (quantified as the distance to word centre in characters. For log-frequency we used the UK or US version of SUBTLEX depending on the corpus and included the log-frequency of the past two words to capture spillover effects.

Note that, while for skipping, we used a *non-linguistic* baseline, for reading times we use a *non-contextual* baseline. This is because for skipping the most interesting contrast is between the role of non-linguistic oculomotor control vs an account that explains skipping via ease of next-word identification (either through prediction or preview). For reading times, by contrast, the most interesting comparison is between properties of the word itself versus contextual cues, as a purely non-linguistic account for gaze duration variation seemed less plausible (indeed, see Figure S12 for a supplementary analysis confirming that the limited relative importance of a purely non-linguistic account for reading time variation).

### Model evaluation

We compared the ability of each model to account for the variation in the data by probing prediction performance in a 10-fold cross-validation scheme, in which we quantified how much of the observed variation in skipping rates and gaze durations could be explained.

For reading times, we did this using the coefficient of determination, defined via the ratio of residual and total sum of squares: 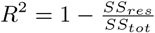. The ratio 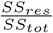 relates the error of the model (*SS*_*res*_) to the error of a ‘null’ model predicting just the mean (*SS*_*tot*_), and gives the variance explained. For skipping, we use a tightly related metric, the McFadden *R*^2^. Like the *R*^2^ it is computed by comparing the error of the model to the error of a null model with only an intercept: 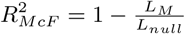, where *L* indicates the loss.

While *R*^2^ and 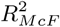 are not identical, they are formally tightly related – critically, both are zero when the prediction is constant (no variation explained) and go towards one proportionally as the error decreases to zero (i.e. towards all variation explained). Note that in a cross-validated setting, both metrics can become negative when prediction of the model is worse than the prediction of a constant null-model.

### Variation partitioning

To assess relative explanatory power, we used variation partitioning to estimate how much of the explained variation could be attributed to each set of explanatory variables. This is also known as *variance* partitioning, as it is originally based on partitioning sums of squares in regression analysis; here we use the more general term ‘variation’ following [78].

Variation partitioning builds on the insight that when two (groups of) explanatory variables (*A* and *B*) both explain some variation in the data *y*, and *A* and *B* are independent, then variation explained by combining *A* and *B* will be approximately additive. By contrast, when *A* and *B* are fully redundant – e.g. when *B* only has an *apparent* effect on *y* through its correlation with *A* – then a model combining *A* and *B* will not explain more than the two alone.

Following [79], we generalise this logic to three (groups of) explanatory variables, by testing each individually and all combinations, and use set theory notation and graphical representation for its simplicity and clarity. For three groups of explanatory variables (*A, B* and *C*), we first evaluate each separately and all combinations, resulting in 7 models:

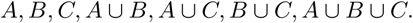

From these 7 models we obtain 7 ‘empirical’ scores (expressing variation explained), from which we derive the 7 ‘theoretical’ variation partitions: 4 overlap partitions and 3 unique partitions. The first overlap partition is the variation explained by all models, which we can derive as:

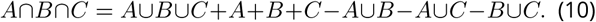

The next three overlap partitions contain all pairwise intersections of models that did not include the other model:

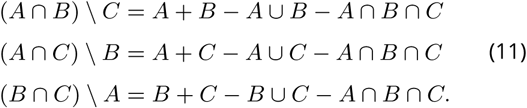

The last three partitions are those explained exclusively by each model. This is the relative complement: the partition unique to *A* is the relative complement of BC: *BC*^*RC*^. For simplicity we also use a star notation, indicating the unique partition of *A* as *A*^*^. These are derived as follows:

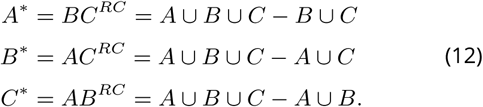

Note that, in the cross-validated setting, the results can become paradoxical and depart from what is possible in classical statistical theory, such as partitioning sums of squares. For instance, due to over-fitting, a model that combines multiple EVs could explain *less* variance than all of the EVs alone, in which case some partitions would become negative. However, following [79], we believe that the advantages of using cross-validation outweigh the risk of potentially paradoxical results in some subjects. Partitioning was carried out for each subject, allowing to statistically assess whether the additional variation explained by a given model was significant. On average, none of the partitions were paradoxical.

### Simulating effect sizes

Preview benefits were simulated as the expected diïňĂerence in gaze duration after a preview of average informativeness versus after no preview at all (…). This this best corresponds to an experiment in which the preceding preview was masked (e.g. XXXX) rather than invalid (see Discussion). To compute this we compared the took the difference in parafoveal entropy between an average preview and the prior entropy. Because we standardised our explanatory variables, this was transformed to subject-specific z-scores and then multiplied by the regression weights to obtain an expected effect size.

For the predictability benefit, we computed the expected difference in gaze duration between ‘high’ and ‘low’ probability words. ‘High’ and ‘low’ was empirically defined based on the human-normed cloze probabilities in provo, which we divided into thirds using percentiles. The resulting cutoff points (low < 0.02; high >0.25) were log-transformed, applied to the surprisal values from GPT-2, and multiplied by the weights to predict effect sizes. Note that these definitions of ‘low’ and ‘high’ may appear low compared to those in literature – however, most studies collect cloze only for specific ‘target’ words in relatively predictable contexts, which biases the definition of ‘low’ vs ‘high’ probability. By contrast, we analysed cloze probabilities for *all* words, yielding these values.

### Statistical testing

Statistical testing was performed across participants within each dataset. Because two of the three corpora had a low number of participants (10 and 14 respectively) we used non-parametric bootstrap t-tests, by creating resampling a null-distribution with zero mean counting how likely a t-value at least as extreme as the true t-value was to occur. Each test used at least 10^4^ bootstraps; p-values were computed without assuming symmetry (equal-tail bootstrap).

## Data and code availability

The Provo and Geco corpora are freely available ([31, 32]). All additional data and code needed to reproduce these results will be made public at the Donders Data Repository.

## Contributions

Conceptualisation: MH. Data wrangling and preprocessing: JvH. Formal analysis: MH, JvH. Statistical analysis and visualisations: MH, JvH. Supervision: FPdL, PH. Initial draft: MH. Final draft: MH, JvH, PH, FPdL.

## Acknowledgements

We thank Maria Barrett, Yunyan Duan, and Benedikt Ehinger for useful input and inspiring discussions during various stages of this project, and Ashley Lewis for helpful comments on an earlier version of this manuscript. This work was supported by The Netherlands Organisation for Scientific Research (NWO Research Talent grant to M.H.; NWO Vidi grant to F.P.d.L.; Gravitation Program Grant Language in Interaction no. 024.001.006 to P.H.) and the European Union Horizon 2020 Program (ERC Starting Grant 678286 to F.P.d.L).

## Supplementary materials

**Figure S1.**
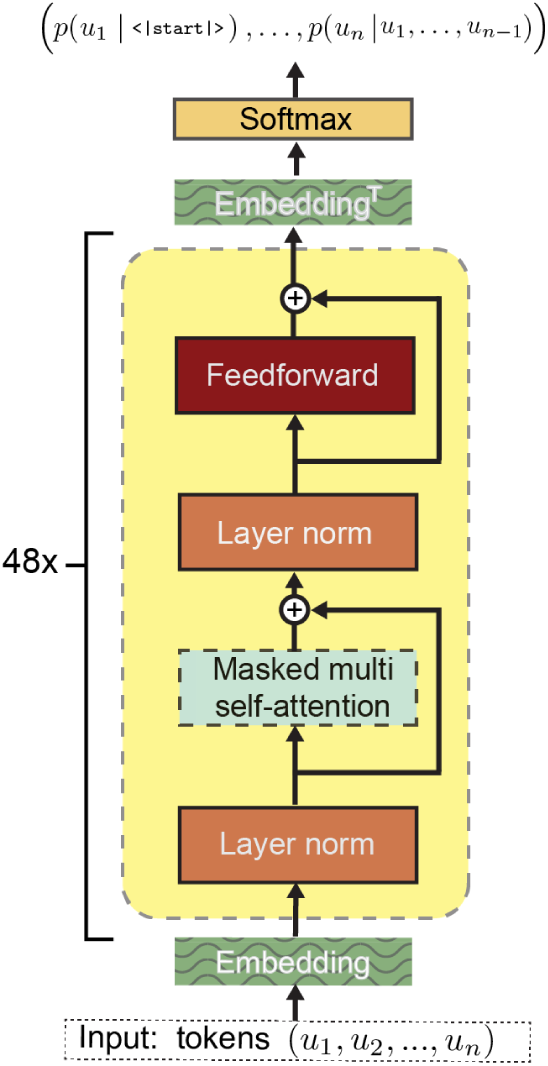
GPT-2 Architecture. Note that this panel is based on the original GPT schematic, with some operations modified and re-arranged to reflect the slightly different architecture of GPT-2. The most important and distinctive step of each transformer block is masked multi-headed self-attention (see Methods). Not visualised here is the initial tokenisation, mapping a sequence of characters into a sequence of tokens.

**Figure S2.**
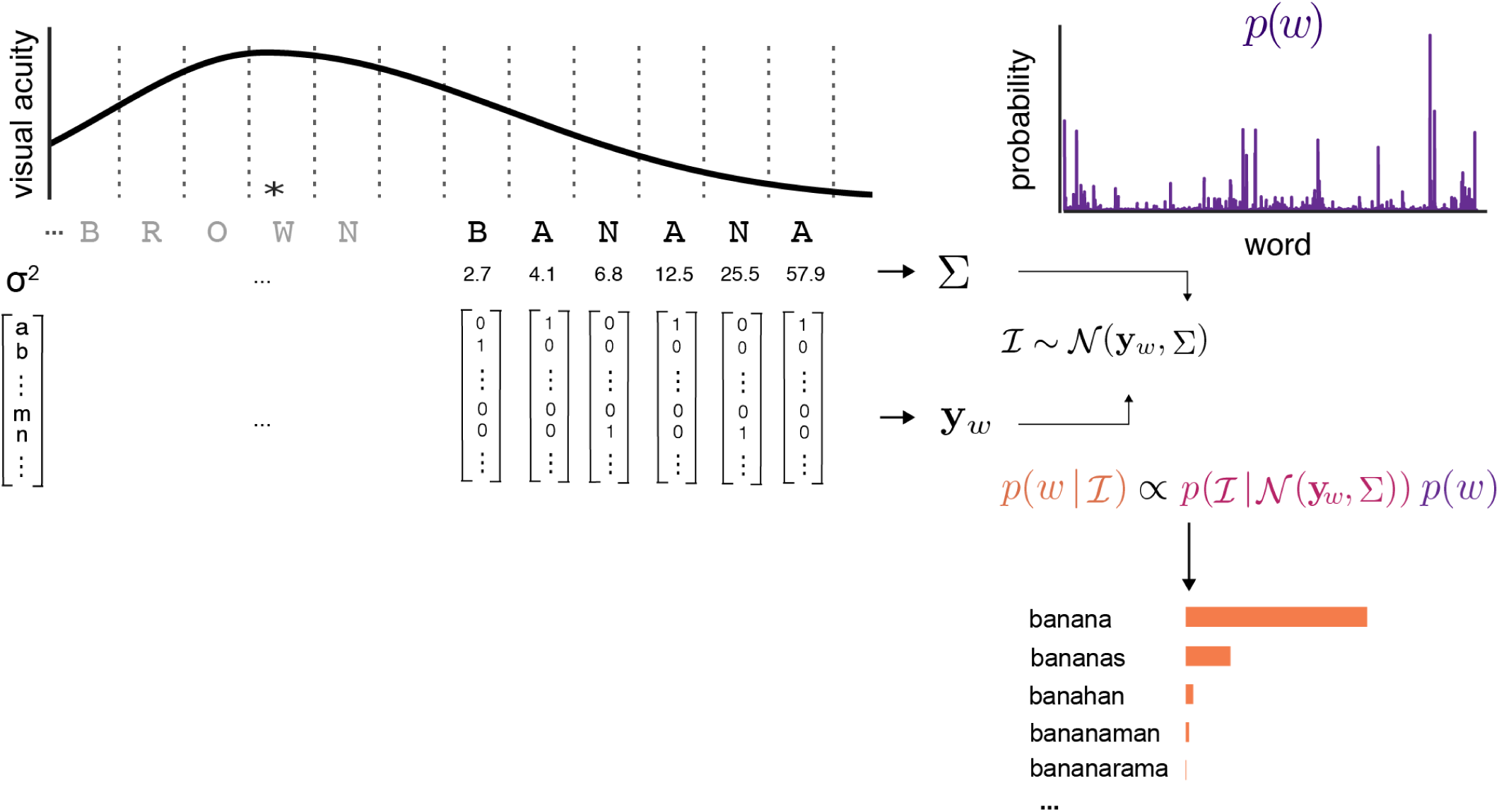
Encoding and inference scheme of the ideal observer analysis. Visualisation of the Ideal Observer, following formulation in [35]. A word at a given eccentricity is converted into a noisy visual percept, after which a posterior probability of the identity of the word given the noisy percept was computed using Bayesian inference. The uncertainty of this posterior (expressed in terms of Shannon entropy) was then used to quantify the expected uncertainty in the parafoveal percept – or, inversely, a word’s *parafoveal identifiability*. In this scheme, words are represented as a concatenation of one-hot encoded letter vectors. Visual information (ℐ) is sampled from a multivariate Gaussian centred on the word vector y_*w*_ with a diagonal covariance matrix Σ, the values of which (*σ*^2^) are inversely related to the integral under the visual acuity function around each letter. The posterior is then computed by comining the likelihood of the visual information ℐ given a particular word, with a prior probability of that word *p*(*w*) (e.g. derived from lexical frequency). This computation was performed using a log-odds formulation that exploits the proportionality in Bayes’ rule to perform belief-updating without renormalisation (see Methods).

**Figure S3.**
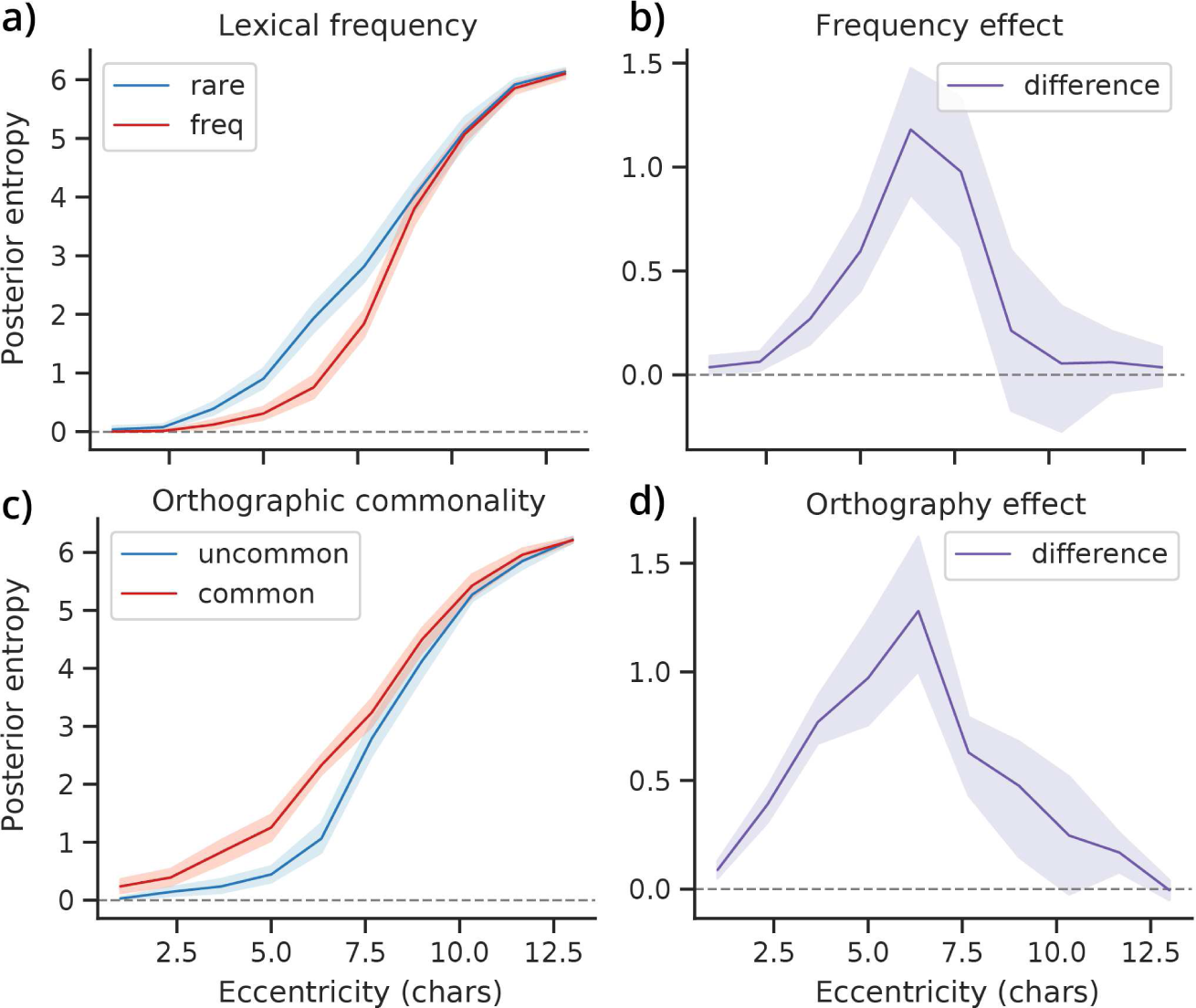
Modulation of parafoveal identifiability by visual and linguistic features, and their interaction. The parafoveal entropy for a given word (Fig S2) is a complex function that integrates linguistic and visual characteristics, and which can account for various known effects, such as the effect of lexical frequency and orthographic neighbourhood on visual word recognition. To illustrate this, we simulated some characteristic effects of eccentricity, frequency (**a**,**b**) and orthographic distinctiveness (**c**,**d**). For frequency (**a**), we randomly sampled 20 ‘rare’ and ‘frequent’ 5-letter words (based on a quartile split), and computed the parafoveal identifiability (quantified via posterior entropy) at increasing eccentricities. As can be seen, the percept becomes uncertain at increasing eccentricities more quickly for low-frequency words, showing that lexical frequency boosts parafoveal identifiability. For orthography (**c**), we similarly sampled 20 7-letter words that were classified as orthographically common or uncommon based on the first three letters. Here, commonality was again defined using a quartile split but now on the number of alternative words starting with the same three letters. For instance, the letters ‘awk’ in the word ‘awkward’ are highly uncommon and allow to identify the entire word with high confidence based on just those three letters. As can be seen, the model predicts that orthographic uniqueness boosts parafoveal identifiability – as observed in experiments (see [13]). Notably, when we consider the difference between the two classes of words (**b**,**d**), an inverted U shape is apparent: the effects are strongest at intermediate visibility. This demonstrates the well-established fact that the effects of prior (linguistic) knowledge is strongest at intermediate levels of perceptual uncertainty (see [42] for discussion). (Note that, while both the orthography and frequency effects are effects of “prior linguistic knowledge”, only the frequency effect is technically an effect of the *prior*, since the orthography effect is driven by the generative model.) In all plots, thick lines represent the mean entropy across words; shaded regions indicate bootstrapped 95% confidence intervals.

**Figure S4.**
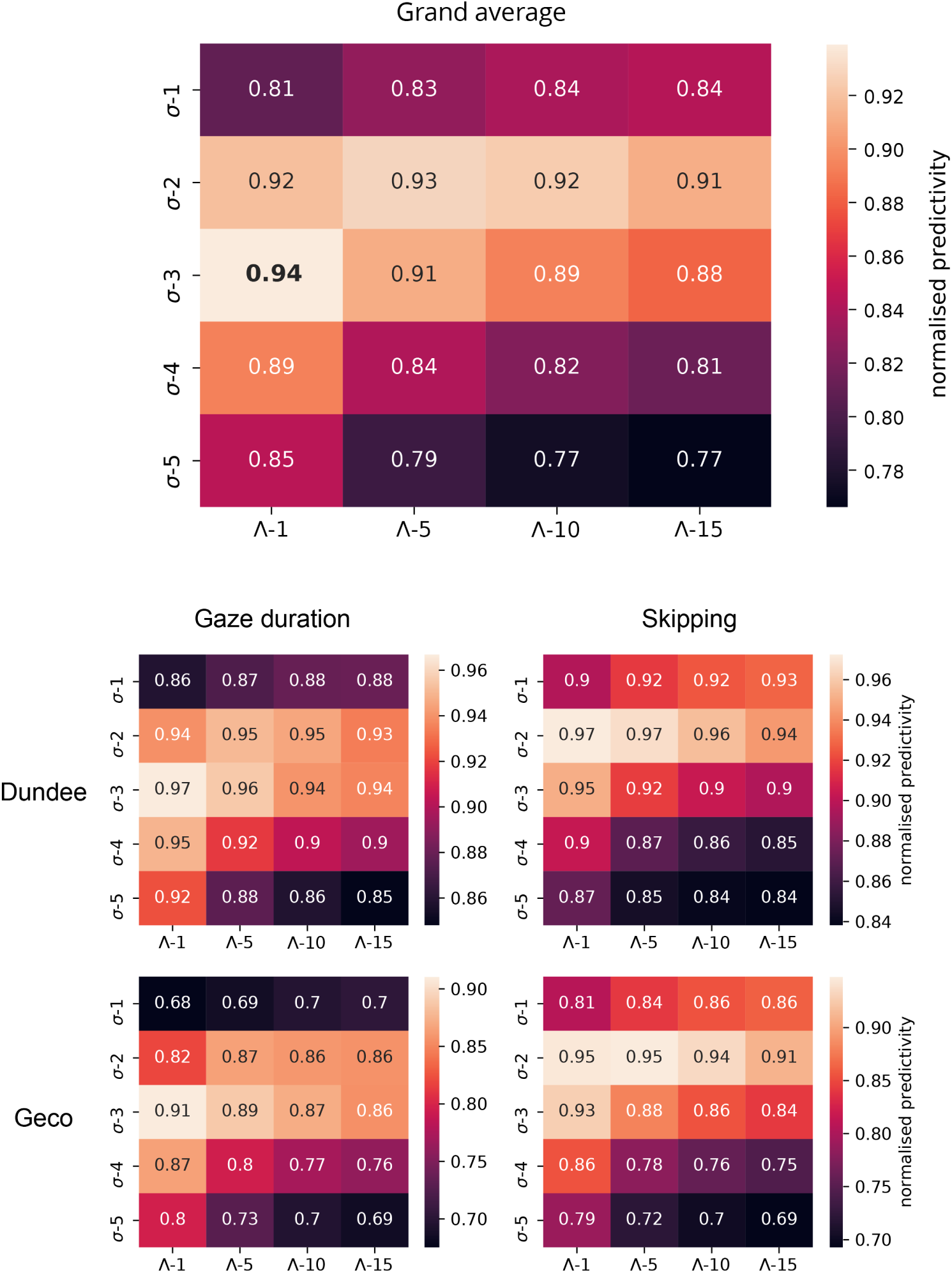
Grid search to establish ideal observer parameters. Grid search result grand average (top) and individual results for different corpora and analyses (bottom). To decide on the values for *σ* and Λ, a grid search was performed on a random subset of 25% of the Dundee and Geco corpus; we did not apply it to PROVO because there was not enough data per participant. In both skipping and reading times, we performed a 10-fold cross-validation with the full model, using parafoveal entropy as computed with different visual acuity parameters *σ* and Λ (Equation 6). To avoid biasing the contextual vs non-contextual model comparison (Figure 5), we used both the contextual and non-contextual prior and averaged the results to obtain the results for each analysis in each corpus. To ensure that different analyses and corpora are weighted equally in the grand average, the prediction scores (*R*^2^ or 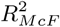) were normalised by dividing the prediction score of each parameter combination by the highest score (i.e. score of the best parameter combination) for each subject, for each analysis. This resulted in *σ* = 3 and Λ = 1, which we have used in all analyses. Note that *σ* determines the perceptual span (see Figure S2) and that *σ* = 3 corresponds well to what is known about the size of the perceptual span and is close to default parameters in other models (see Methods).

**Figure S5.**
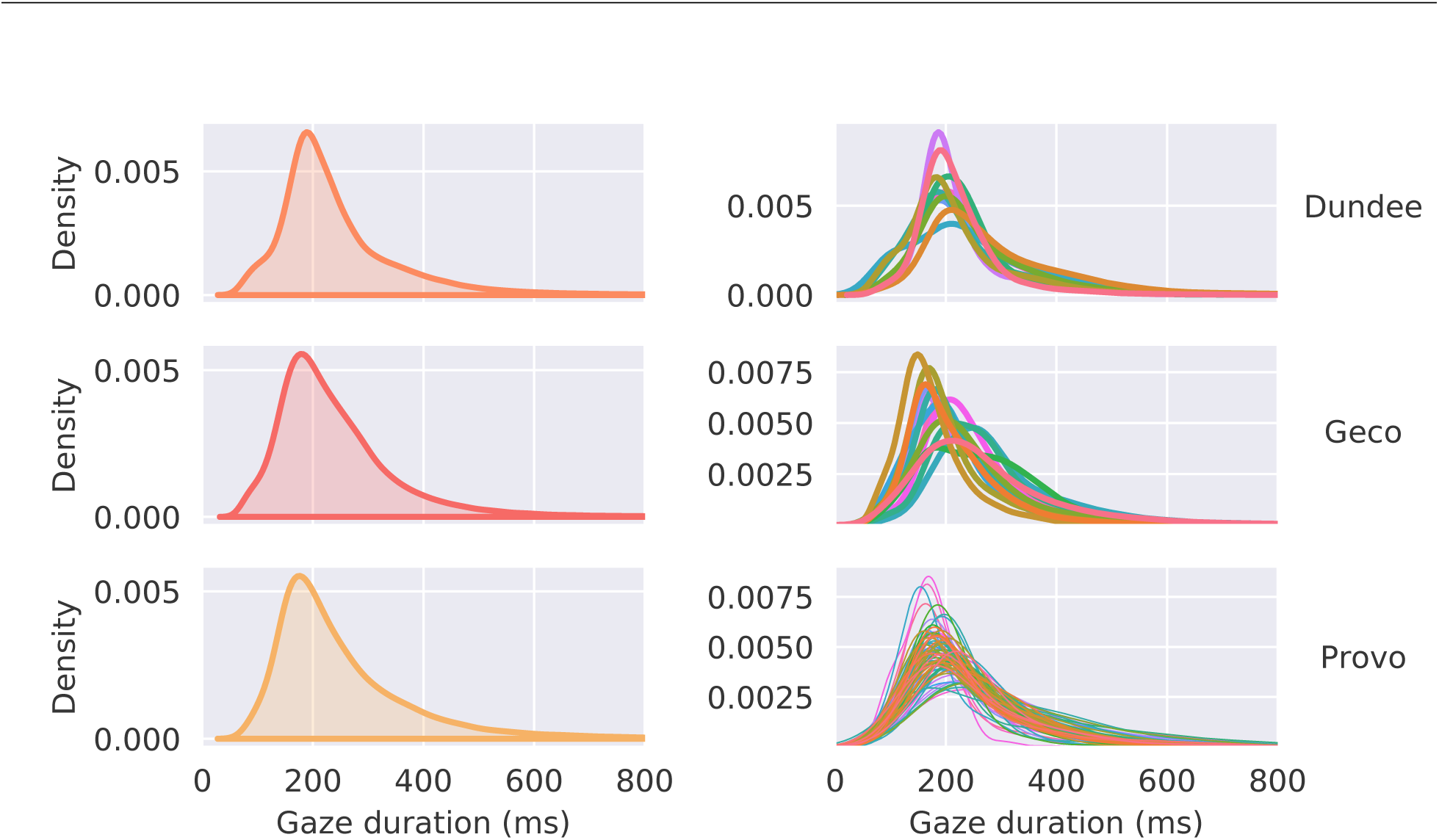
Distributions of reading times (gaze durations). Kernel density estimate of the distribution of reading times across all datasets, both on average (left column) and in individual participants (right column).

**Figure S6.**
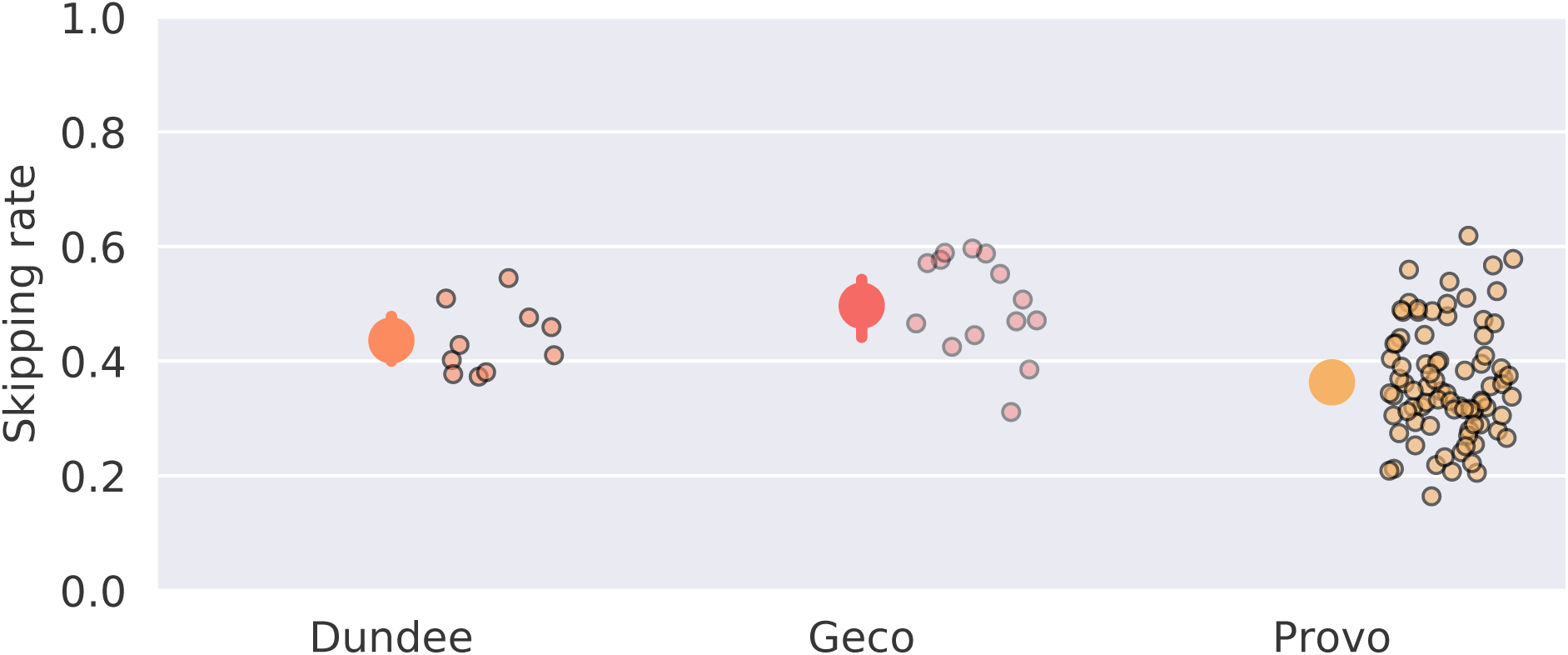
Average skipping rate in each dataset. Average rate of skipping in all words included in the skipping analysis (see *Methods*) in all datasets. Large dots with error bar show group mean plus bootstrapped 95% confidence interval. Small dots show indidividual participants.

**Figure S7.**
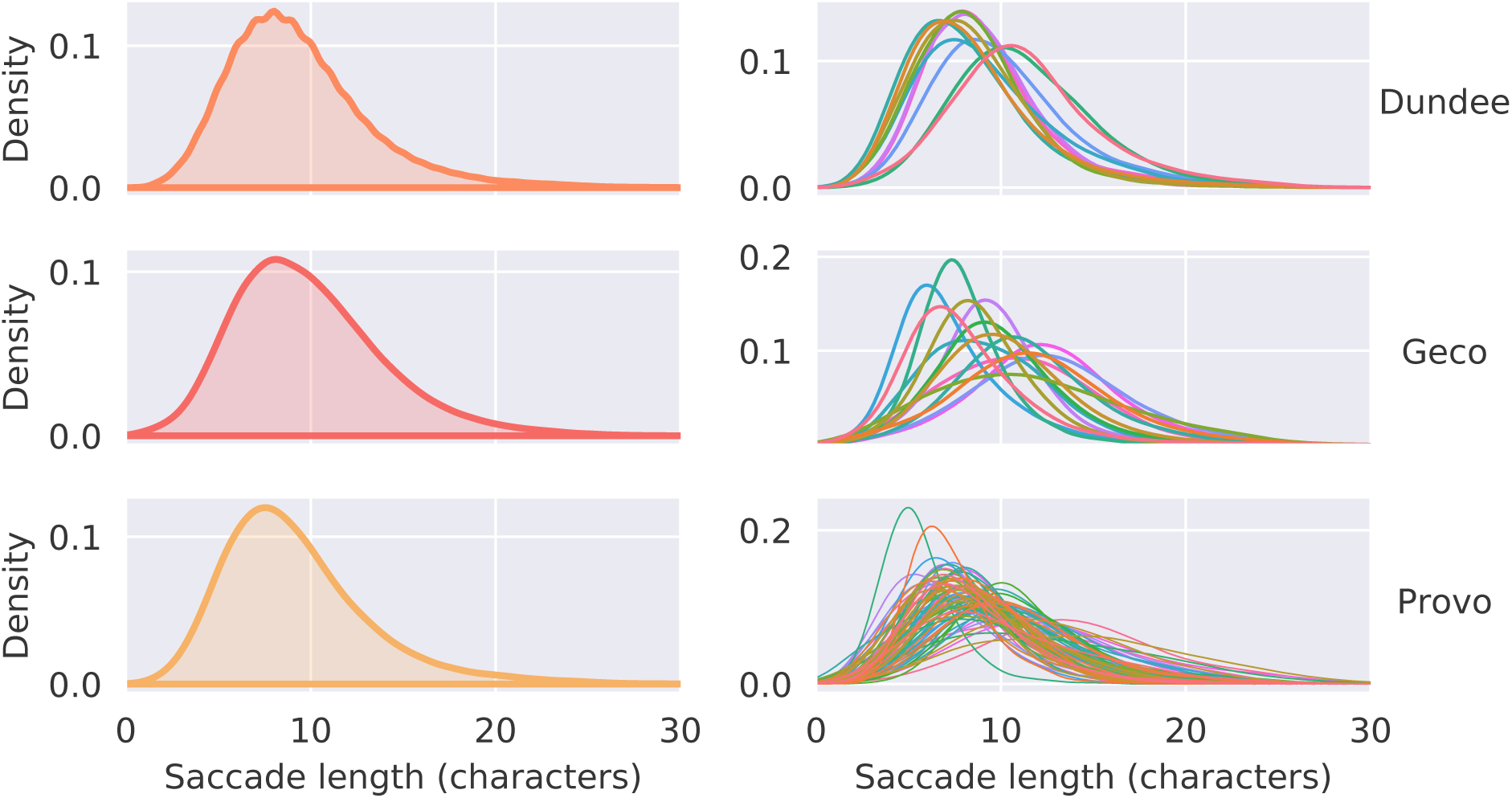
Distribution of (forward) saccade lengths in each datasets. Kernel density estimate of the distribution of saccade lengths (amplitudes) of first-pass, forward saccades in all datasets, both on average (left column) and per individual participant (right column). Note that for this visualisation we only included progressive, forward saccades within the same line (excluding saccades that cross lines), up to a maximum amplitude of 24 characters (excluding saccades during periods participants were not actually reading).

**Figure S8.**
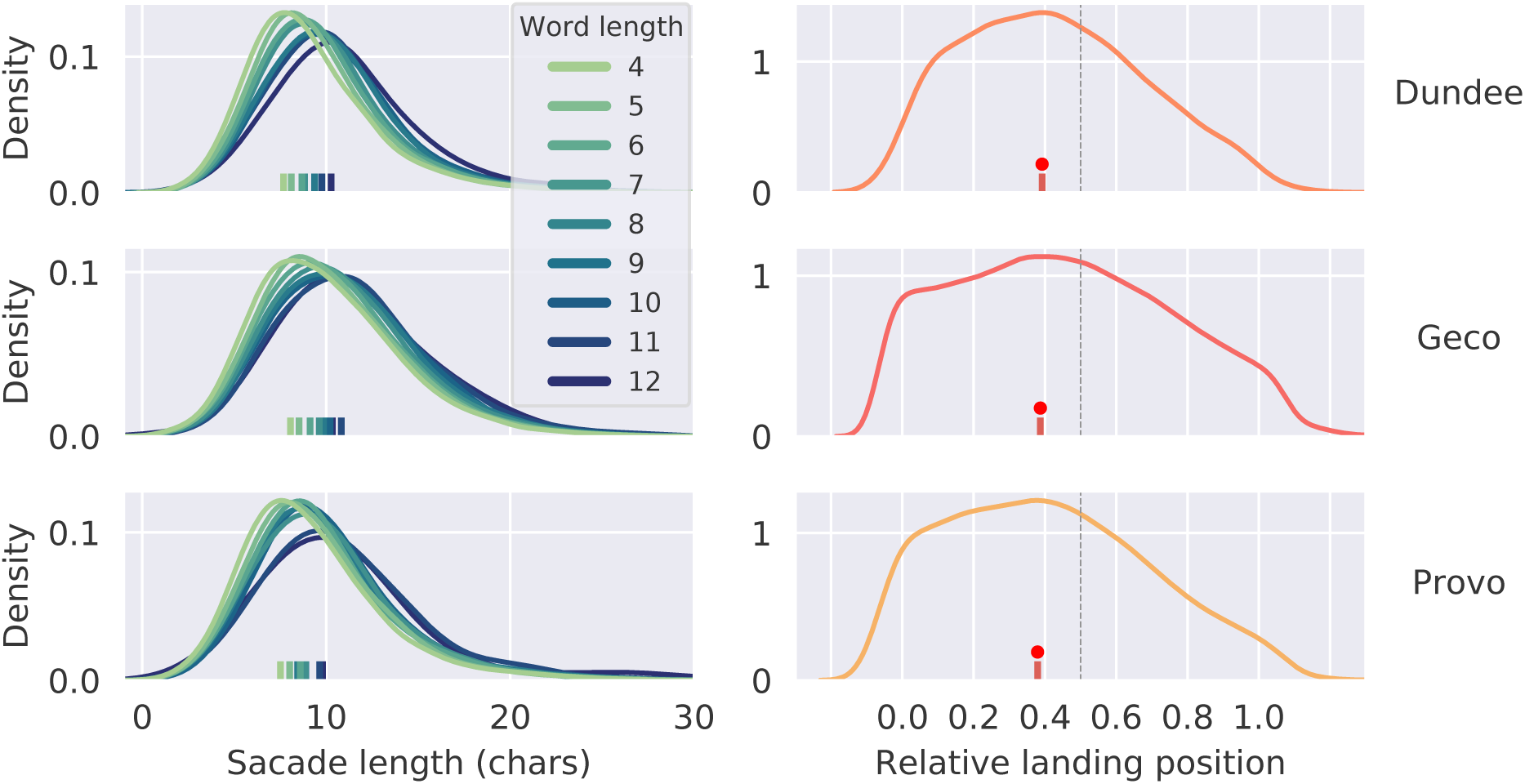
Saccade lengths are tailored to word lengths and exhibit a preferred landing position. **Left column**: Kernel density estimate of the saccade lengths, estimated separately for target words of different lengths. Colours indicate word lengths, vertical lines indicate the mode of the distribution. **Right column**: Kernel density estimate (plus mode) of the relative landing position, averaged across words with different lengths. Saccades are longer for longer words, such that a systematic ‘preferred landing position’ is maintained, slightly left to the center of the word (indicated by the vertical dashed line); see [45, 46].

**Figure S9.**
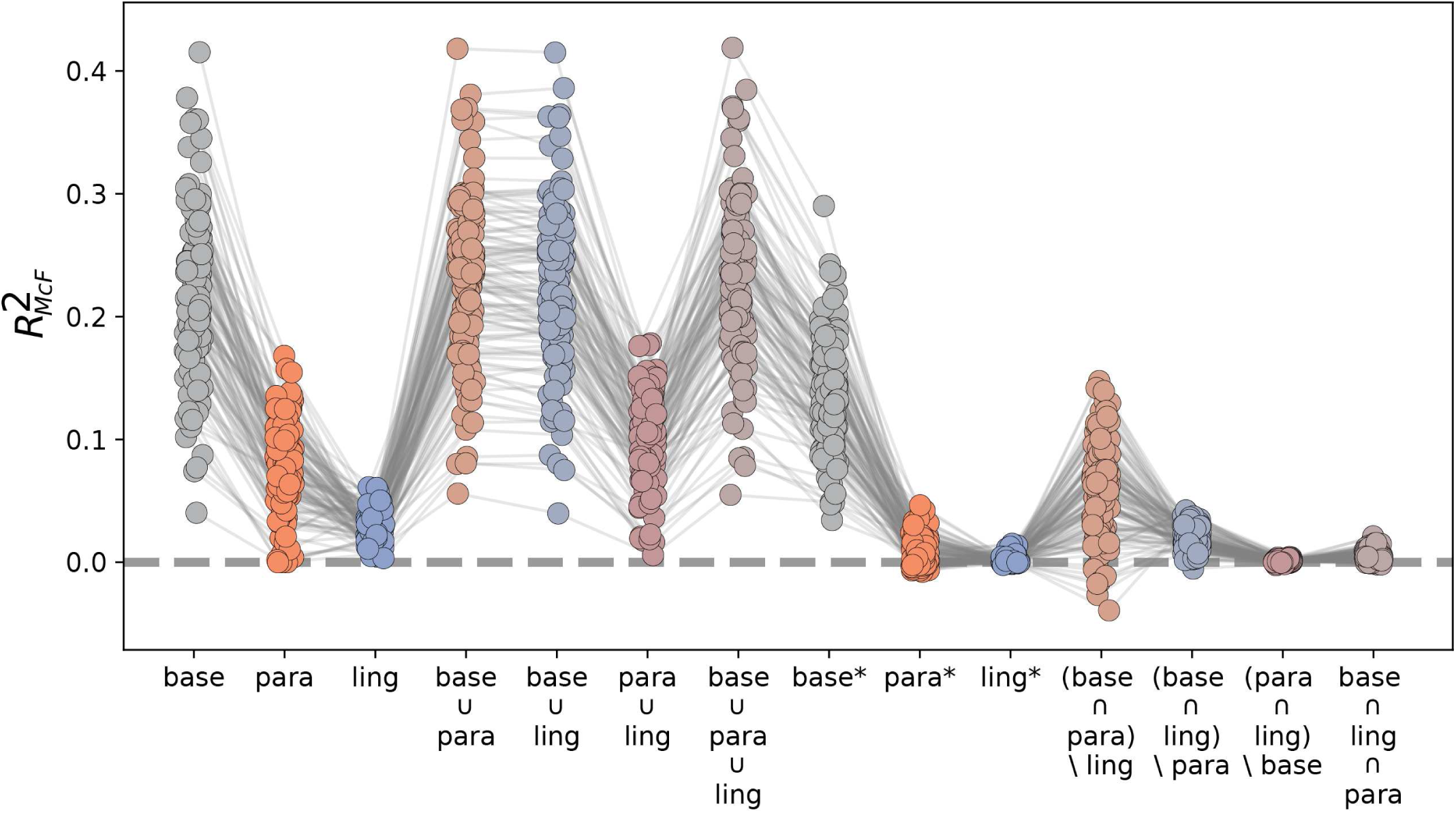
Skipping variation partitioning for all participants. Explained cross-validated variation partition for skipping (see Fig 2) of each partition, for each participant, for the skipping analysis. Models for the baseline, parafoveal preview and linguistic prediction are indicated by ‘base’, ‘para’, and ‘ling’, respectively. Unions are indicated by ∪, intersections by ∩; for the relative complement we use the asterisk-notation: e.g. ‘para*’ indicates variation explained uniquely by parafoveal preview. Note that due to cross-validation, the amount of variation explained can become negative in some partitions for individual participants (see Methods).

**Figure S10.**
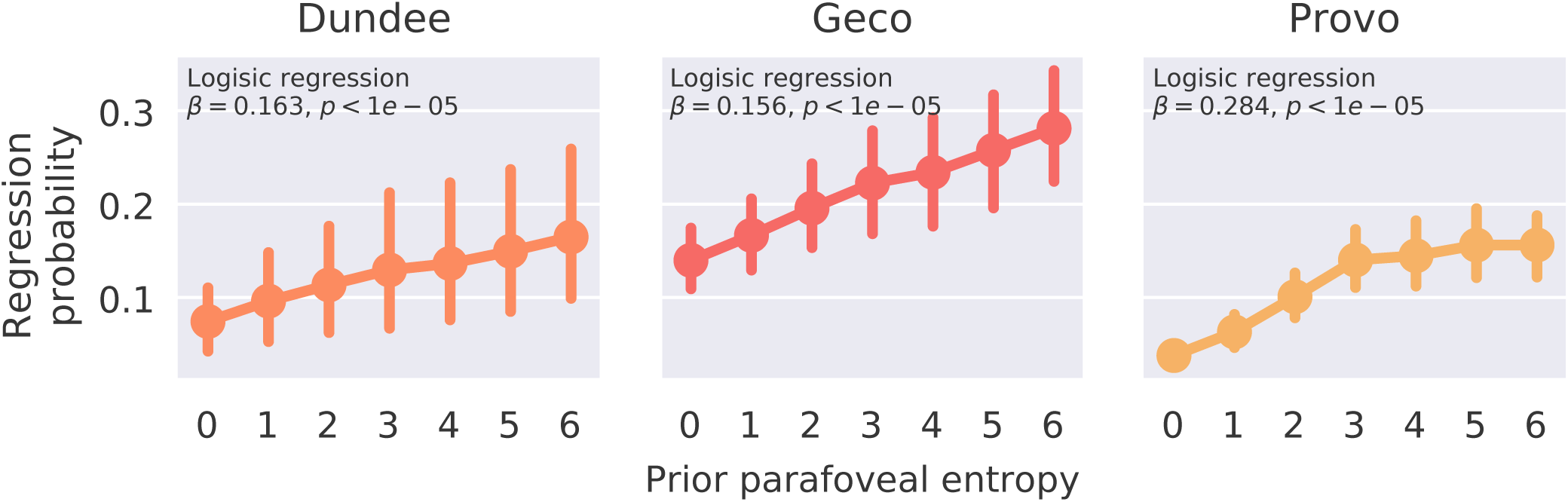
Probability that a skipped word is regressed to depends on its prior identifiability. Probability that an initially skipped word is subsequently fixated (i.e. regressed to), as a function of the prior parafoveal entropy, before skipping. Dots with connecting lines show the average regression probability for initially skipped words as a function of the binned prior parafoveal entropy. Error bars show the (bootstrapped) 95% confidence interval around the mean (across participants). In all datasets, the probability that a skipped word gets subsequently fixated depends on the amount of visual information about word identity that was available *before* the word was was skipped, suggesting a compensation mechanism. Note that the binning is done for visualisation purposes only. Statistical evaluation is based on a subject-wise logistic regression on the word-by-word parafoveal entropy and regression values. Statistical significance is established by a bootstrap test on the subjects’ coefficients, in each dataset.

**Figure S11.**
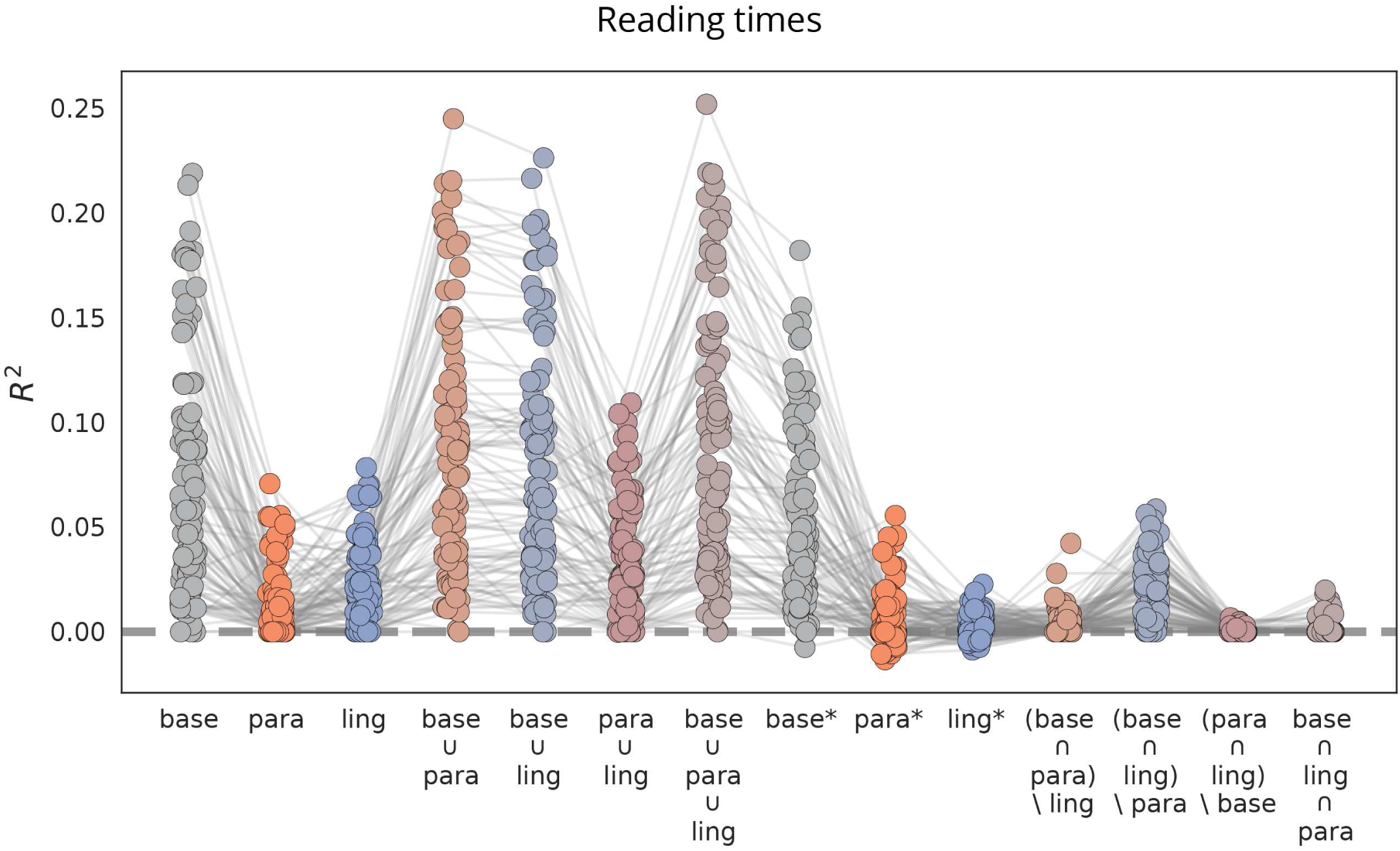
Reading times variance partitioning. Explained cross-validated variation partition for skipping (see Fig 3) of each partition, for each participant, for the skipping analysis. Models for the baseline, parafoveal preview and linguistic prediction are indicated by ‘base’, ‘para’, and ‘ling’, respectively. Unions are indicated by ∪, intersections by ∩; for the relative complement we use the asterisk-notation: e.g. ‘para*’ indicates variation explained uniquely by parafoveal preview (see Methods). Note that due to cross-validation, the amount of variation explained can become negative in individual participants (see Methods).

**Figure S12.**
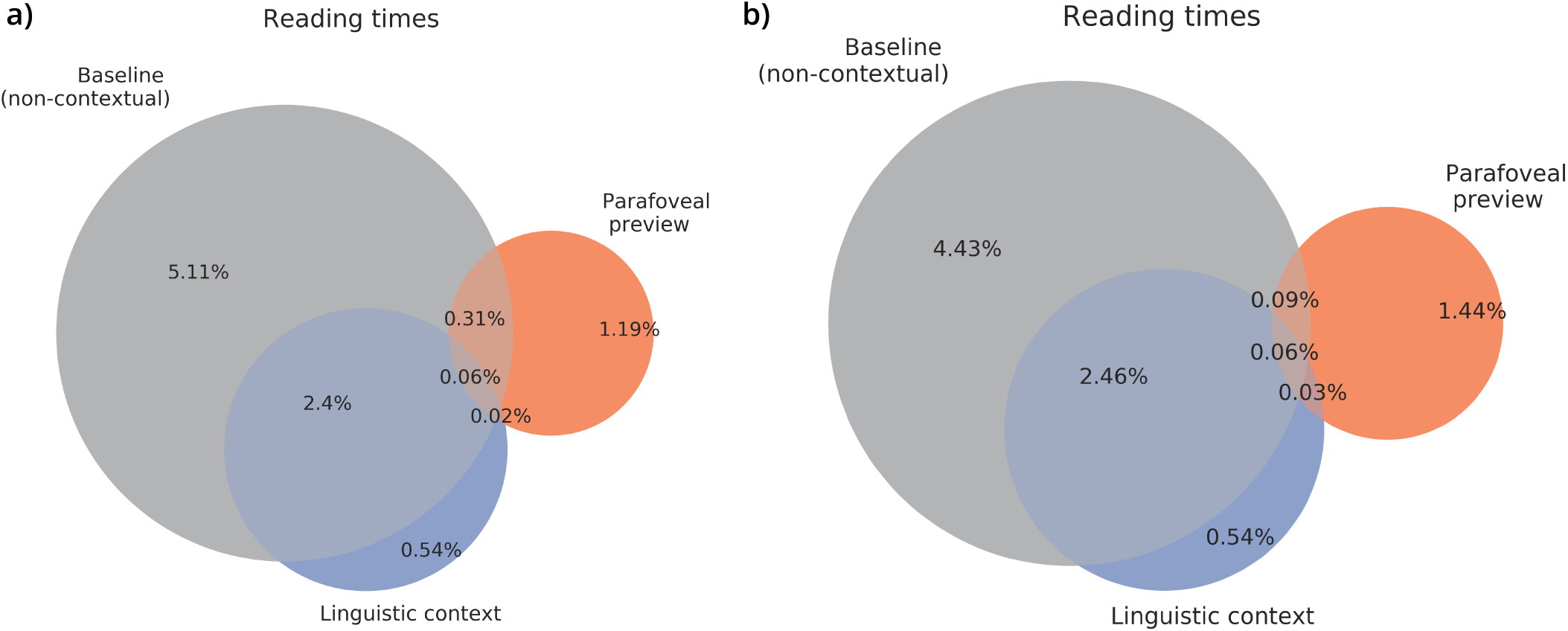
Reading times variance partitioning with and without non-linguistic factors. Same as in Fig 3, but comparing the baseline with (**a**)) or without (**b**)) the primary non-linguistic explanatory factor for reading time variation – viewing position [80]. Including the viewing position adds 0.7% additional variance explained. This demonstrates while that viewing position affect reading times, the amount of variance uniquely explained by non-linguistic factors is much lower for reading times than for skipping.

**Table S1.** Literature sample for effect size ranges

| Effect type | Publication | Effect size |
| --- | --- | --- |
| preview benefit | Inhoff, A. W. (1989). Lexical access during eye fixations in reading: Are word access codes used to integrate lexical information across interword fixations?. *Journal of Memory and Language*, 28(4), 444-461. | 51 |
| preview benefit | Veldre, A., & Andrews, S. (2018). Parafoveal preview effects depend on both preview plausibility and target predictability. *Lexical access during eye fixations in reading: Quarterly Journal of Experimental Psychology*, 71(1), 64-74. | 49 |
| preview benefit | Inhoff, A. W., & Rayner, K. (1986). Parafoveal word processing during eye fixations in reading: Effects of word frequency. *Perception & psychophysics*, 40(6), 431-439. | 40 |
| preview benefit | McDonald, S. A. (2006). Parafoveal preview benefit in reading is only obtained from the saccade goal. *Vision Research*, 46(26), 4416-4424. | 35 |
| preview benefit | Williams, C. C., Perea, M., Pollatsek, A., & Rayner, K. (2006). Previewing the neighborhood: The role of orthographic neighbors as parafoveal previews in reading. *Journal of Experimental Psychology: Human Perception and Performance*, 32(4), 1072. | 26.7 |
| preview benefit | Kennison, S. M., & Clifton, C. (1995). Determinants of parafoveal preview benefit in high and low working memory capacity readers: Implications for eye movement control. *Journal of Experimental Psychology: Learning, Memory, and Cognition*, 21(1), 68. | 25.25 |
| preview benefit | Blanchard, Harry E., Alexander Pollatsek, and Keith Rayner. "The acquisition of parafoveal word information in reading." *Perception & Psychophysics* 46.1 (1989): 85-94. | 22.6 |
| preview benefit | Schroyens, W., Vitu, F., Brysbaert, M., & d'Ydewalle, G. (1999). Eye movement control during reading: Foveal load and parafoveal processing. *The Quarterly Journal of Experimental Psychology Section A*, 52(4), 1021-1046. | 14.6 |
| prediction benefit | Ehrlich, S. F., & Rayner, K. (1981). Contextual effects on word perception and eye movements during reading. *Journal of verbal learning and verbal behavior*, 20(6), 641-655. | 33 |

Table S1 – *Continued from previous page*
| Effect type | Publication | Effect size |
| --- | --- | --- |
| prediction benefit | Rayner, K., & Well, A. D. (1996). Effects of contextual constraint on eye movements in reading: A further examination. *Psychonomic Bulletin & Review*, 3(4), 504-509. | 20 |
| prediction benefit | RJ. Altarriba, J. Kroll, A. Sholl, K. Rayner. (1996) The influence of lexical and conceptual constraints on reading mixed-language sentences: Evidence from eye fixations and naming times *Memory & Cognition*, 24 (1996), pp. 477-492. | 21 |
| prediction benefit | Ashby, J., Rayner, K., & Clifton Jr, C. (2005). Eye movements of highly skilled and average readers: Differential effects of frequency and predictability. *The Quarterly Journal of Experimental Psychology Section A*, 58(6), 1065-1086. | 23.5 |
| prediction benefit | Rayner, K., Ashby, J., Pollatsek, A., & Reichle, E. D. (2004). The effects of frequency and predictability on eye fixations in reading: implications for the EZ Reader model. *Journal of Experimental Psychology: Human Perception and Performance*, 30(4), 72 | 19 |
| prediction benefit | Rayner, K., Binder, K. S., Ashby, J., & Pollatsek, A. (2001). Eye movement control in reading: Word predictability has little influence on initial landing positions in words. *Vision Research*, 41(7), 943-954. | 15 |
| prediction benefit | Rayner, K., Slattery, T. J., Drieghe, D., & Liversedge, S. P. (2011). Eye movements and word skipping during reading: effects of word length and predictability. *Vision Research*, 41(7), 943-954. | 18 |
| prediction benefit | Hand, C. J., Miellet, S., O'Donnell, P. J., & Sereno, S. C. (2010). The frequency-predictability interaction in reading: It depends where you're coming from. *Journal of Experimental Psychology: Human Perception and Performance*, 36(5), 1294-1313. | 12 |

## Notes

### Competing Interest Statement

The authors have declared no competing interest.

### Summary of Updates

Improved discussion. Improved information on underlying eye movement data and acquisition. Additional analysis of regressions as compensation mechanism for low-level saccade target selection.

